# Non-equilibrium modeling of flux through biomolecular condensates

**DOI:** 10.64898/2026.02.03.703649

**Authors:** Yashraj M. Wani, Jerelle A. Joseph

**Affiliations:** Department of Chemical and Biological Engineering, Princeton University, Princeton, NJ, USA; Omenn–Darling Bioengineering Institute, Princeton University, Princeton, NJ, USA

## Abstract

Cellular biomolecular condensates function far from equilibrium, sustained by continuous influx and efflux of molecular components. Notable examples include the nucleolus, P-bodies, and stress granules, in which regulated molecular transport is essential for their function. Experimental characterization of molecular flux through condensates remains challenging due to limitations in spatiotemporal resolution, making computational approaches a powerful alternative for systematic investigation. Here, we present a computational approach (TRACE) to model molecular transport through biomolecular condensates under non-equilibrium steady-state conditions at residue-level resolution. Using TRACE, we systematically probe physicochemical factors that govern molecular flux through model condensates. We find that protein sequence composition and patterning determine the internal ‘mesh size’ of the condensates, which strongly dictates molecular transport efficiency, and the ballistic-to-diffusive dynamics crossover of the fluxing molecules. We also find that molecule stiffness can enhance their flux by exposing larger surface for intermolecular interactions. Furthermore, we find that modulating host-guest interactions can modulate solubility and friction which jointly governs molecular flux. Finally, we use our framework to show that crosslinking reduces condensate pore size, hindering molecular flux – consistent with experimental observations of dextran diffusion through nucleolar phases. Collectively, these insights provide a basis for rationally tuning condensate-mediated molecular flux, with potential applications in therapeutics and soft material design.

## I. INTRODUCTION

Over the past decade, numerous experimental studies have provided compelling evidence that membraneless organelles in cells form through phase separation of proteins, RNA and DNA.^1–4^ The resulting biomolecular condensates serve as key organizers of intracellular biochemistry, participating in a wide range of essential functions including gene expression, stress response, and metabolism.^1,5^ A hallmark of condensates in living cells is that they are inherently out-of-equilibrium, with a constant exchange/flux of material with their surrounding environment, driven by biochemical activity such as ATP hydrolysis and enzymatic reactions, among others.^6,7^ Notable examples include the nucleolus, where ribosomal RNA is synthesized and processed as it fluxes through distinct phases;^8^ stress granules and P-bodies, where RNA-binding proteins and RNAs shuttle in and out continually;^9–11^ and the nuclear pore complex, where intrinsically disordered FG-nucleoporins form a selective barrier controlling molecular transport between the nucleus and cytoplasm.^12,13^

Despite the ubiquity and importance of molecular exchange, the factors governing molecular fluxes through condensates remain poorly understood, for several reasons. Fluorescence recovery after photobleaching (FRAP), the most common technique for probing condensate dynamics, measures the ensemble-averaged behavior of labeled molecules, where the recovery signal can reflect several processes that are difficult to decouple, including diffusion within the condensate, exchange across the interface, and binding/unbinding kinetics.^14,15^ Setting this aside, inferring diffusion coefficient by bleaching a small region within a large condensate can be highly sensitive to the choice of underlying mathematical model.^14^ Finally, fluorescent tags can alter the interactions and mobility of the probe molecules, skewing the observations.^16–18^ Higher resolution can be achieved with single particle tracking (SPT) where molecules are sparsely labeled so that their individual trajectories can be reconstructed and analyzed.^19–21^ SPT, however, requires specialized microscopy setups and also suffers from the issue of fluorescence tags affecting the observations. Additionally, confinement of labeled molecules within the condensate can lead to poor sampling of the interfacial and dilute phases in conventional SPT, requiring specialized algorithms for proper sampling of the dilute phase.^22^ Overall, due to technical limitations, experimental studies probing biomolecular condensates out-of-equilibrium are mostly qualitative.

In contrast, computational methods such as molecular dynamics (MD) simulations offer the required spatiotemporal precision and control to probe the physics of flux through condensates. For instance, molecular simulations have significantly advanced our understanding of the physical principles underlying biomolecular phase separation – helping explain the thermodynamics, structure, and dynamics of biomolecular condensates.^23–35^ Through coarsegrained modeling,^30,36–38^ and atomistic simulations,^26,39–41^ researchers have demonstrated how specific molecular interactions – such as hydrophobic and charged interactions – contribute to phase separation and under what environmental conditions (e.g. pH, salt, temperature) condensates form. These studies have also provided insights into the internal organization (such as multiphasic architecture) and material properties (e.g., surface tension and viscosity) of condensates.^40,42–45^

While these simulations have been effective in revealing the equilibrium/near-equilibrium physics of condensates, they capture only part of the story, as many natural condensates exist out of equilibrium. Accordingly, there is a growing need to design computational frameworks to model non-equilibrium phenomena in biomolecular condensates.^15,46^ Recent works include molecular dynamics coupled with Metropolis Monte Carlo method to model non-equilibrium dynamics of protein phosphorylation, and chemically informed modifications to force fields to investigate the non-equilibrium liquid to solid transition in condensates.^47,48^

In this work, we aim to systematically characterize molecular flux through condensates and to identify the physical principles that influence it. Understanding the physics of transport within condensates is a critical step toward a broader framework in which condensates are understood not simply as phase-separated compartments, but programmable bioreactors whose throughput can be predicted, tuned, and ultimately engineered. To achieve this goal, we introduce the TRACE (‘TRansport Across Condensate Environments’) approach to model non-equilibrium steady-state molecular flux through condensates with residue-level resolution. We use TRACE to probe flux of glycine based peptides through model protein condensate composed of tyrosine and serine residues and then probe how crosslinking of condensate proteins affects molecular flux.

Our results demonstrate that sequence composition and residue patterning determine the internal architecture of condensates, which in turn largely control transport properties. We further show that molecule stiffness facilitates flux through condensates, and that associative interactions between condensate and fluxing molecules modulate their solubility and diffusivity, jointly determining the net flux turnover. Finally, we employ TRACE to probe the effect of crosslinking on molecular flux, finding that crosslinking shrinks condensate pore sizes and thereby hinders flux – consistent with previous experimental interpretations.

## II. RESULTS

### A. An approach to capture sequence-dependent molecular flux through condensates

We seek to investigate the flux of biomolecules through condensates, motivated by cellular structures such as the nucleolus. Transcriptional machinery localized at the core of the nucleolus continuously synthesizes rRNA molecules, which exhibit radially outward flux as they undergo processing to ultimately leave the nucleolus as pre-ribosomal subunits.^8,50^ Molecular flux arises as a consequence of biochemical processes that consume energy.

To simulate molecular flux, we choose a coarse-grained resolution that allows us to describe a condensate comprising of at least a few hundred proteins and to probe flux over hundreds of nanoseconds, which can be prohibitively expensive in an all-atom resolution. In this work, we focus primarily on protein-based condensates, while anticipating that the physical principles uncovered will be broadly applicable across diverse condensate systems.

In coarse-grained simulations, reactions – such as energy consuming processes – have been accounted for implicitly using various approaches that include probabilistic topology updates,^47^ or using specialized interactions using virtual sites.^51^ In that spirit, to probe molecular flux and elucidate the underlying physicochemical mechanisms, we develop a framework (TRACE) that explicitly captures the two essential components for studying molecular flux: (i) an internal ‘active site’ where underlying biochemical activities constantly produce material – referred henceforth as *guest molecules* – generating a concentration gradient that results in an entropic driving force on the guest molecules and (ii) a surrounding *host condensate* that mediates their transport and eventual release. From a technical perspective, our objective is to develop an approach capable of capturing both sequence and interaction dependent effects that may modulate molecular flux in condensates.

To capture the sequence-dependent effects we adopt a residue-resolution coarse-grained model, Mpipi, where each amino acid is represented as a single bead and beads interact with each other in a chemically specific manner.^36^ Importantly, the Mpipi model captures the critical solution temperatures of intrinsically disordered proteins (IDPs) in agreement with experiments, making it an appropriate choice in this study.^52^ Here, we represent both the host and the guest molecules via the Mpipi model, while the active site is represented semi-implicitly as discussed below.

Direct-coexistence simulations have been successfully employed to study the phase behavior of IDPs at thermodynamic equilibrium [Fig. S1(a)].^25,30,34,36,53,54^ In direct-coexistence simulations, the condensed and dilute phases are simulated in the same simulation cell and from such simulations one can compute phase diagrams, critical temperatures, interfacial tensions, key interactions that drive phase separation of IDPs, among other properties.^25,40,45,55–57^ Starting from a direct-coexistence simulation – with the condensate as a spherical droplet or modeled in the slab geometry^30^ – we introduce an *active site* at the center of the ‘host’ condensate using a discrete particle surface comprising of particles arranged on a lattice, with their identities chosen to mimic that of the host protein.^58^ Fig. 1(a) shows a spherical(left) and planar(right) active sites within the protein condensates. The construction of the active site is discussed in greater detail in the methods section. Functionally, chemical composition of the active site promotes wetting of the condensate, preventing any drift of the host condensate.

**FIG. 1:**
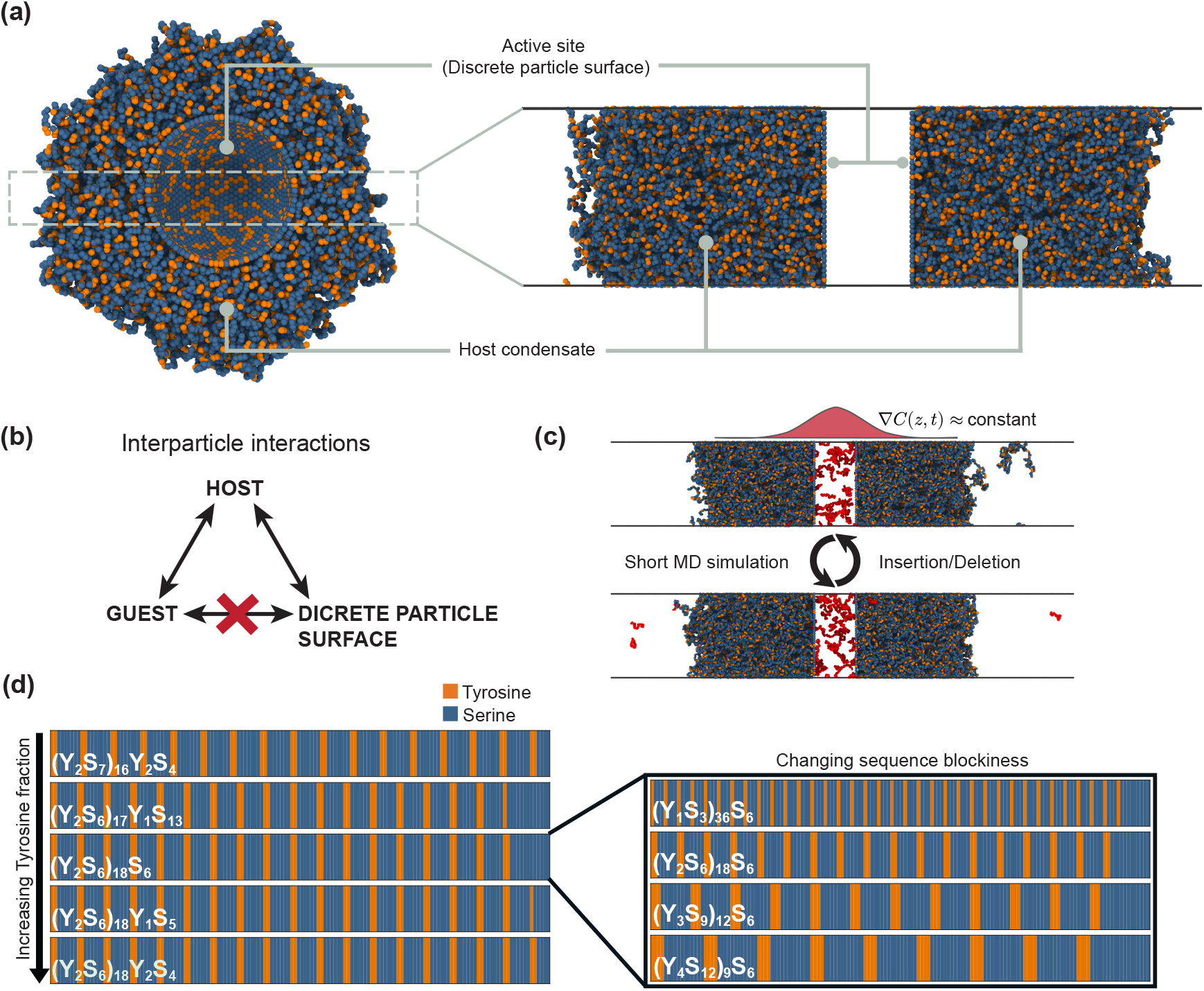
Simulation framework to probe flux through biomolecular condensates. (a) Simulation setup showing a spherical active site wetted by a protein condensate (left) and a slab approximation in the low-curvature limit (right), with host condensate flanking the active site on both sides. The active site is described by a semi-permeable discrete particle surface with the same amino acid composition as the host. Simulation snapshots rendered with Ovito-pro visualization software.^49^ (b) Semi-permeability of the active site implemented through selective interparticle interactions that permit guest permeation while excluding host molecules. (c) Hybrid molecular dynamics scheme to maintain steady flux of guest molecules through the host condensate, consisting of alternating short MD simulations and particle insertion/deletion steps. (d) Schematics of host sequences (150 amino acids) composed of Tyrosine (Y) and Serine (S). Left: increasing tyrosine fraction from top to bottom. Right: increasing sequence blockiness from top to bottom at fixed tyrosine fraction.

As noted earlier, the active site serves as the region where guest molecules are ‘produced’, generating a concentration gradient. These molecules then diffuse outwards, inducing a flux through the host condensate into the surrounding dilute phase. However, a complication arises: interactions between the guest molecules and the active site surface act a boundary, effectively trapping guest molecules. To resolve this, we parameterize the interparticle interactions such that the active site surface is essentially ‘invisible’ to the guest molecules, ensuring that the guest molecules can freely exit the active site region, enabling unhindered flux [Fig. 1(b)]. In practice, this is achieved by setting *ε*_*ij*_ = 0 in Eq.5 for interactions between active site and guest residues [Fig. S3]. The final framework more closely mirrors what occurs within biomolecular condensates, where there is no distinct boundary between the active site and the surrounding condensate.

Finally, maintaining a steady-state flux requires a constant concentration gradient. To achieve this, we implement a hybrid molecular dynamics scheme [Fig. 1(c)]: the system is first evolved through standard molecular dynamics for a short finite duration, after which an insertion/deletion step is performed. Any guest molecule that has diffused into the dilute phase is removed and reinserted inside the active site. By repeating this two-step cycle multiple times, we maintain an effective and stable concentration gradient throughout the simulation, allowing us to observe and quantify steady-state flux through the condensate over time as would be expected in a real system.

### B. Model host–guest systems to decode principles of molecular flux through condensates

Within the computational framework described above, we can systematically investigate factors that influence molecular flux through condensates. There are several key parameters that can be probed, ranging from the properties of the host condensate, the guest molecules, and the surrounding environment.

The first set of variables concerns the host condensate itself. The sequence composition and architecture of the host proteins directly determine the condensate’s thermodynamic and material properties, including dense-phase density, critical temperature, surface tension, and viscosity.^45,57^ By varying the amino acid sequence, we can modulate the strength and distribution of interactions, thereby altering condensate stability. Additionally, condensate architecture can be adjusted to represent either monophasic or multiphase assemblies,^42^ allowing us to probe how internal organization affects molecular transport.

The second set of variables involves the guest molecules which may represent small peptides, RNAs, or small molecules, and their physicochemical properties can be systematically varied. In the case of polymeric guests such as peptides or RNA, we can tune their intrinsic properties – such as length, residue type, and backbone rigidity – to explore how molecular flexibility and interaction strength influence their flux. More complex topologies, such as proteins containing folded domains or globular regions, can also be introduced to capture behavior relevant to cellular proteins.

Finally, the environmental conditions – including temperature, pH, and molecular crowding – provide another layer of control. While environmental factors can strongly influence condensation and molecular transport, in this work we primarily hold them constant to ensure that observed changes in flux can be attributed to intrinsic molecular and structural features rather than global environmental variables.

Given the wide parameter space, we focus on a minimal system designed to isolate the essential physics of flux through condensates [Fig. 1(d), Fig. S4]. The host condensate consists of disordered proteins, 150 amino acids in length, composed of tyrosine (Y) and serine (S) residues, where tyrosine acts as a hydrophobic residue promoting inter-/intrachain interactions and serines serve as flexible polar linkers. The guest molecules are glycine (G) peptides of varying lengths (10-, 20-, 30-mers), which interact weakly with the host (since the host sequences are predominantly composed of serines) and provide a means to modulate host– guest interactions without perturbing host–host interactions [Fig. S4(b)]. Importantly, this minimal system does not have any charged species which allows us to isolate factors affecting flux in a systematic manner with mainly hydrophobic interactions between residues.

To explore the role of sequence composition and patterning in the host condensate, we adopt two complementary strategies [Fig. 1(d)]. In the first, we vary the overall tyrosine fraction, ranging between 22%–26% tyrosine, thereby tuning the strength of intermolecular interactions [Table S1]. Tyrosine fractions are restricted to a narrow range to ensure that the critical temperatures of the proteins are within the same range (approximately 350 K) and the simulations are performed close to criticality and physiological temperature (310 K) [Fig. S5]. In the second, we fix the composition but rearrange the sequence patterning (blockiness) of residues along the host proteins to assess the role of interaction heterogeneity [Table S2]. At low blockiness, the tyrosine residues are well separated with multiple serines between them, while at high sequence blockiness, multiple tyrosines are clustered together [Fig. 1(d) right]. Sequence blockiness is defined based on the number of Y-S bonds in the sequence [See Methods].

### C. Molecular flux is highly correlated to pore size distributions of host condensates

A key quantity used to characterize flux in our simulations is the average flux time, defined as the time required for a guest molecule to traverse the entirety of the host condensate. The methodology for identifying traversal events and computing flux times is described in detail in the Methods section.

To probe how changes in hydrophobicity affect the average flux time, we systematically varied the tyrosine fractions of the host sequences. We observed that the average flux time increased linearly with increasing tyrosine fraction [Fig. 2(a)], reflecting the concomitant increase in the host condensate density [Fig. S5], which reduces the available pore space and hinders guest transport. In addition, for any given tyrosine fraction of the host proteins, longer guests consistently required more time to traverse the condensate, consistent with their larger radii of gyration and reduced diffusivity [Fig. S11]. Notably, the slope of the linear dependence of flux time on tyrosine fraction scaled linearly with guest lengths [Fig. 2(a) inset], indicating that increasing guest length not only increases the average flux time but also amplifies the sensitivity of transport to changes in host condensate hydrophobicity.

**FIG. 2:**
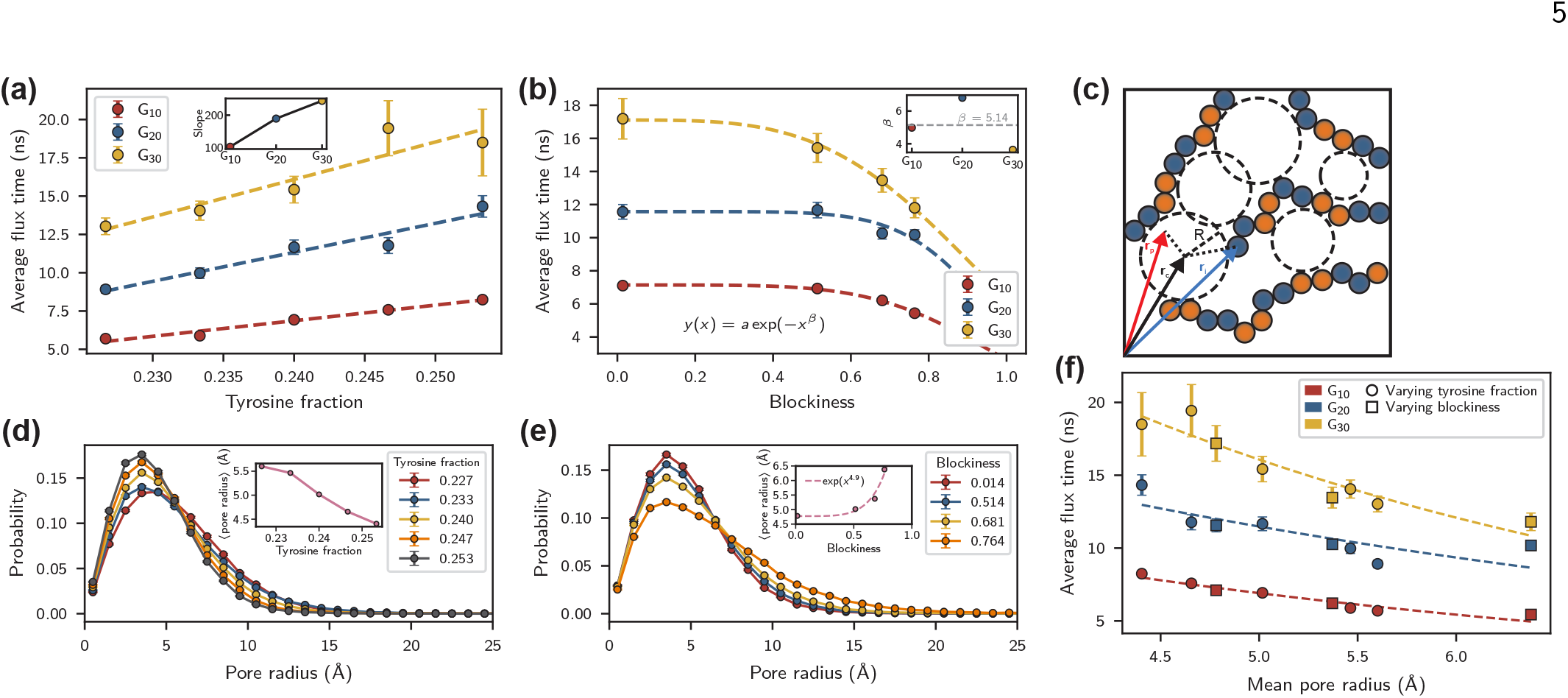
Pore size distribution of host condensates governs guest molecular flux. (a, b) Average flux times for glycine guests of lengths 10 (red), 20 (blue), and 30 (yellow) through host condensates with (a) increasing tyrosine fraction and (b) increasing sequence blockiness. Insets show the slopes and scaling exponents for the fitted curves as a function of guest length, respectively. (c) Two-dimensional schematic illustrating the calculation of the pore size distribution by fitting spheres of varying sizes inside condensate voids such that each sphere just contacts surrounding particles. (d, e) Corresponding pore size distributions for host condensates with (d) varying tyrosine fraction and (e) varying sequence blockiness. Insets show the mean pore radius as a function of tyrosine fraction and blockiness, respectively. (f) Average flux time for all the host–guest combinations as a function of mean pore radius.

Orthogonally, when we varied residue patterning of the host sequences while keeping the overall tyrosine fraction identical, the flux times exhibited a non-linear decay with increasing host sequence blockiness [Fig. 2(b)]. This relationship was well captured by a stretched exponential decay function. This behavior can be attributed to the emergence of structural heterogeneity within the condensate: as blockiness increases, tyrosine-rich regions form dense subdomains, while serine-rich regions remain relatively dilute.^56,57^ This spatial inhomogeneity may create uneven pore sizes for guest molecules to flux through, leading to the observed non-linear dependence.

To validate these interpretations, we examined the pore size distributions of the host condensates by fitting spheres in the void spaces of the host condensate (refer to Methods section), conceptualizing them as dynamic meshes through which guest molecules diffuse [Fig. 2(c)]. In general, shape of the pore size distributions resemble that of skewed normal distributions. Increasing the tyrosine fraction shifted the distribution toward smaller pore sizes, with the mean pore size decreasing linearly with tyrosine fraction – consistent with denser packing and slower flux [Fig. 2(d)], in line with the linear dependence observed in Fig. 2(a). In contrast, increasing sequence blockiness broadened the distribution and produced a sharp increase in the mean pore size, reflecting the emergence of dense and sparse regions characteristic of structurally heterogeneous condensates [Fig. 2(e)], consistent with the non-linear dependence observed in Fig. 2(b). Interestingly, we see that both the mean pore size and the average flux time can be fit to a stretched exponential as a function of sequence blockiness, with the stretching exponent in the ballpark for both cases.

Notably, plotting the average flux time against mean pore radius collapsed all observations onto a common trend line [Fig. 2(f)], reinforcing that structural homogeneity or heterogeneity of the host condensate plays a key role in regulating the flux of guest molecules. Flux through condensates depends not only on the total number of attractive interactions in the host sequences, but also on how these interactions are arranged along the sequence. When attractive interactions are more separated and not clustered – giving rise to a more homogeneous architecture – molecular flux is slowed. Conversely, when attractive interactions are bunched together, producing a more inhomogeneous architecture, molecular flux is enhanced.

### D. Host condensates act as poor solvents for guest molecules, which exhibit ballistic-to-diffusive transport crossover at time and length scales corresponding to the pore size of the host condensates

Next, we analyzed the conformational properties of guest molecules as they flux through the host condensates. In Fig. 3(a) we plot the spatial variation in radius of gyration of the guest molecules, along the long axis of the simulation box. We observed that the guest molecules were slightly compact within the condensate relative to their dilute phase dimensions, indicating that the condensate acts as a poor solvent for glycine guests. The overall magnitude of compaction was modest (up to 5%), suggesting that, for the peptides studied, conformational restriction contributes only minimally to the observed flux behavior. To assess whether this compaction was a direct consequence of molecular flux or an inherent property of the condensate environment, we performed NVT (constant molecule count, volume and temperature) simulations of the dense phase of the host condensates mixed with a small number of guest molecules (discussed in detail in the Methods section). These simulations revealed a similar degree of guest molecule compaction, confirming that the observed conformational changes arise from the environment that the host condensate presents to the guests rather than from the flux process [Fig. S6, S15].

**FIG. 3:**
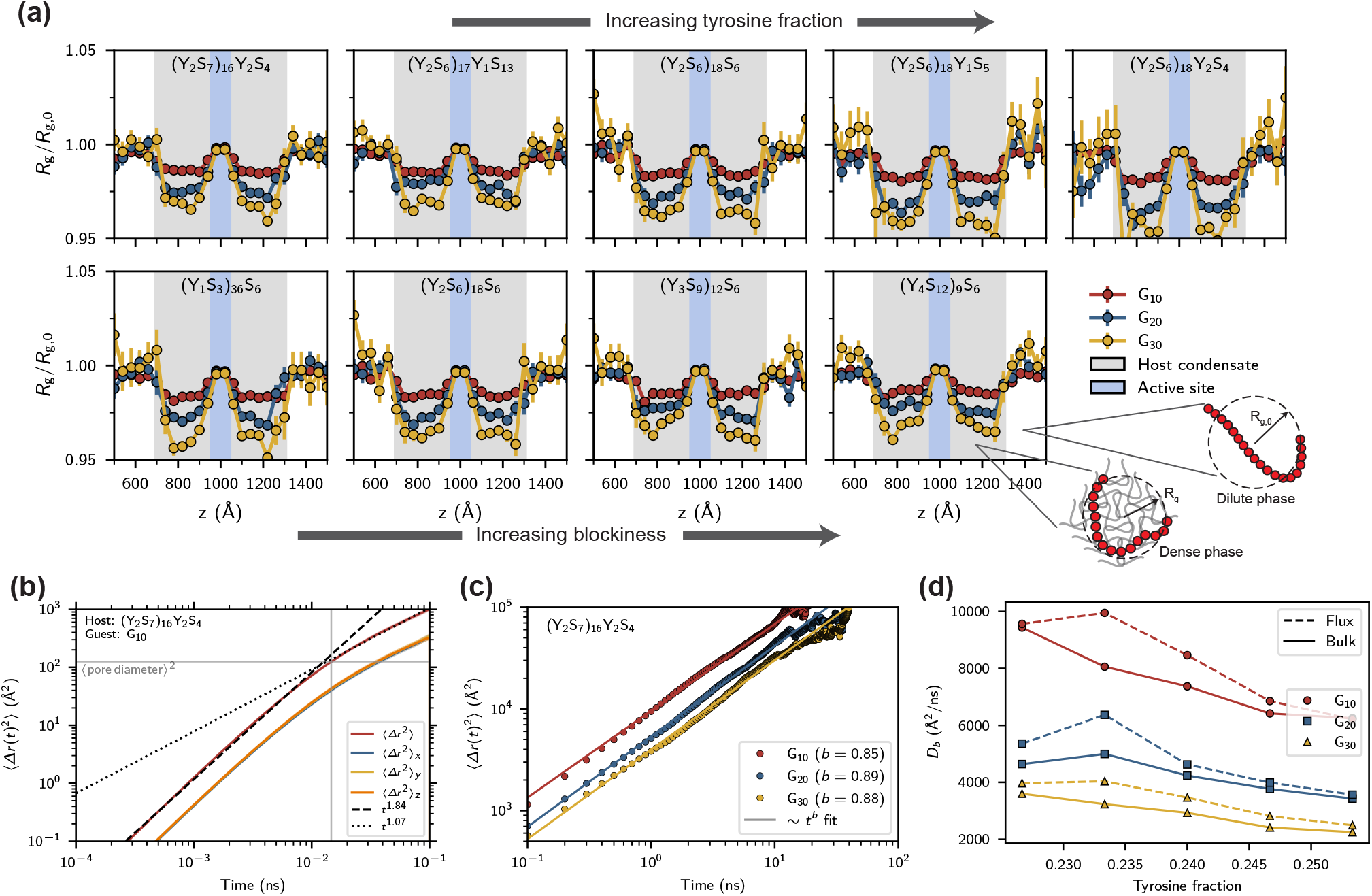
Host condensate acts as a poor solvent for glycine guests, where they exhibit ballistic dynamics within local cages defined by condensate pore sizes and diffusive dynamics at longer times. (a) Spatial variation of the guest radius of gyration relative to dilute phase value *R*_g,0_, as guest molecules traverse the host condensates. Gray and blue color patches indicate the host condensate and active site regions, respectively. Top: increasing tyrosine fraction (left to right). Bottom: increasing sequence blockiness (left to right). (b) Mean squared displacement of guest molecules (G_10_) during transport through the (Y_2_S_7_)_16_Y_2_S_4_ host condensate at short times (*t <* 0.1 ns), with the dashed and dotted lines showing the ballistic and diffusive regimes, respectively. (c) Mean squared displacement of guest molecules during transport through the (Y_2_S_7_)_16_Y_2_S_4_ host condensate at long times (*t >* 0.1 ns). Different colors represent different guest molecules. The solid lines are guide to the eye indicating diffusive transport with scaling exponent ∼0.9. (d) Generalized diffusion coefficient of guest molecules under equilibrium (bulk) and non-equilibrium conditions (flux).

We also analyzed the transport behavior of the guest molecules when inside the host condensates by measuring their center of mass mean squared displacements. At short times (*t <* 0.1 ns) the guests exhibited a crossover between ballistic and diffusive transport of the guest molecules as they move through the host condensates [Fig. 3(b)]. Interestingly we found the crossover occurring at length scales corresponding to the pore diameters of the host condensates, for all the host-guest combinations considered in this study [Fig. S7]. The approximate ballistic motion before the crossover time suggests an unhindered motion of the guest molecules inside a local cage formed by the condensate pores, which becomes diffusive once the guest molecules encounter the host condensate molecules and hop between different local cages. We confirmed that the crossover occurred well before the expected crossover time of ≈ 0.1 ns from the Langevin equations of motion [Fig. S8].

At longer times (*t >* 0.1 ns) the guest molecules, while inside the host condensates, exhibited near diffusive dynamics [Fig. 3(c), Fig. S9], with the MSD scaling exponents between 0.75 and 0.9. As expected, the smaller guest molecules diffused faster than the longer guests through the condensates. Using the proportionality constant for the MSD curves to define generalized diffusion coefficient, *D*_*b*_ = *Δr* (*t*)^2^*/t*^*b*^, we observed a trend of decreasing diffusion coefficient in response to increasing tyrosine fraction in the host sequences [Fig. 3(d)]. In addition, to analyze whether the concentration gradient had any effect on the guest diffusion, we measured the diffusion coefficients from the bulk (NVT) simulations. We observed that the guest molecules consistently diffused slower in the bulk simulations.

### E. Stiffer guest molecules flux faster through host condensates

Biopolymers in nature exhibit diverse topologies and stiffnesses based on their underlying sequence and identity, which can significantly impact their dynamics through condensates. For example, folded domains in proteins such as alpha helices or base pairing in RNAs can render them stiff with persistence lengths on the order of tens of nanometers. To interrogate the effect of stiffness of the guest molecules on their flux, we introduced bending rigidity in the guest molecules by using a harmonic style angle potential,

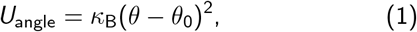

where *κ*_B_ is the strength of the angle potential that determines the chain stiffness and *θ*_0_ = 180^*°*^ is the equilibrium angle. Here, we used *κ*_B_ = 0.0, 1.5, 3.0 kcal/mol/rad^2^, such that at the largest bending stiffness, the guest molecules consisting of 10 monomers become rod-like, as observed from the persistence length calculations [Fig. S11]. As a consequence of the bending stiffness, the radius of gyration approximately doubled for all the guests [Fig. S12], implying an increase in the effective size of the guest molecules.

First, to investigate if the host condensate environment affects the conformations of the guests, we measured the spatial dependence of the guest radius of gyration as they fluxed through the host condensate. From Fig. 4(a), we observe that the stiff guests do not experience conformational changes while traversing through the host condensate, as opposed to the flexible guests that experience compaction when inside the host condensate. For conciseness, in Fig. 4(a) we only show the extremes of the host condensates (i.e., in terms of host hydrophobicity and blockiness), while the remaining radius of gyration profiles are shown in Fig. S13 and Fig. S14 for varying tyrosine fractions and sequence blockiness, respectively. In every case stiff guests do not experience compaction.

**FIG. 4:**
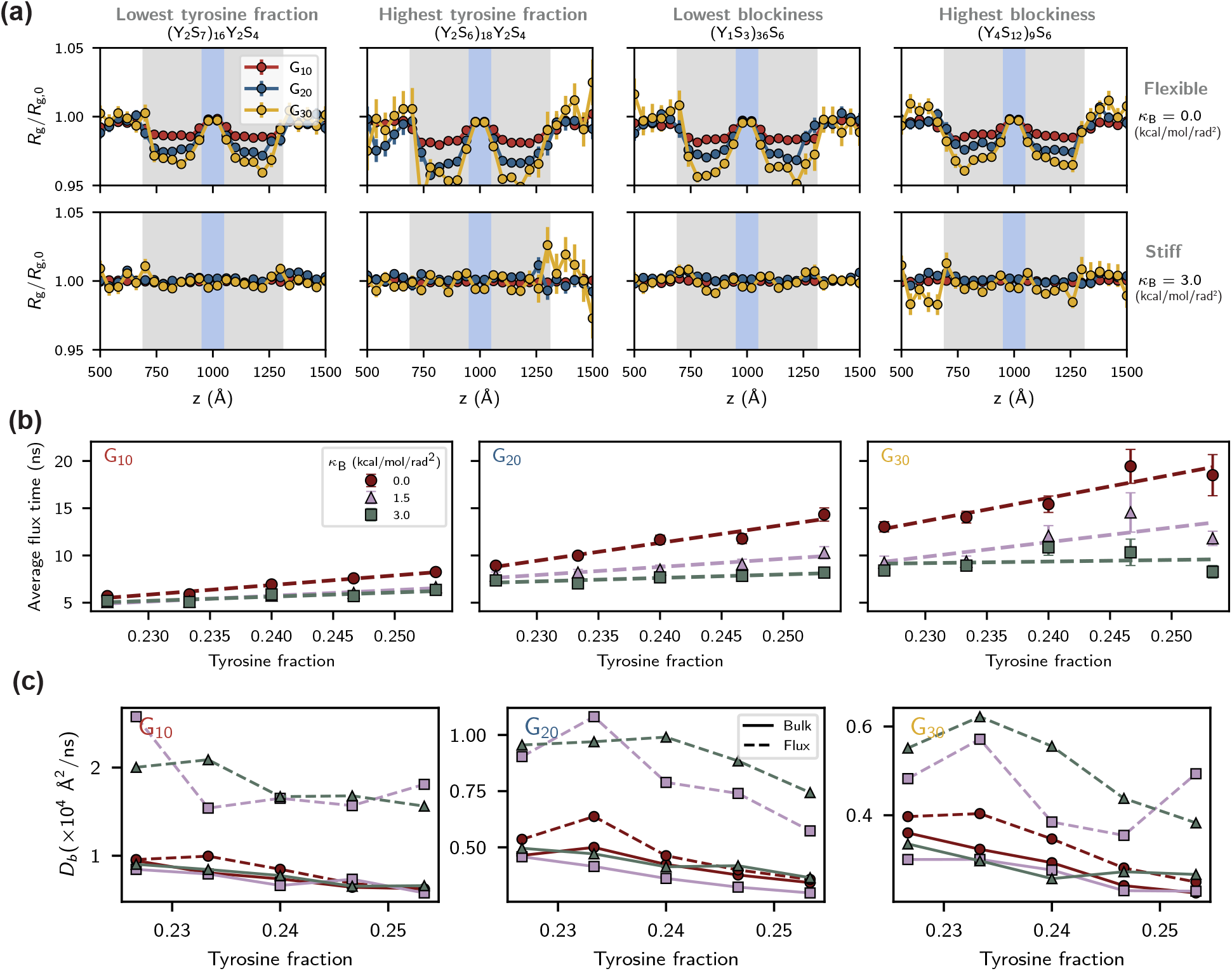
Stiffer guest molecules flux faster through host condensates. (a) Spatial variation of the guest radius of gyration along the long axis of the simulation box. Top: flexible guests (*κ*_B_ = 0.0); Bottom row: stiffest guests (*κ*_B_ = 3.0). Columns 1-2: lowest and highest host tyrosine fraction. Columns 3-4: least and most blocky host sequences. (b) Average flux time as a function of host tyrosine fraction, with different guest molecules – left: G_10_, middle: G_20_, and right: G_30_. (c) Generalized diffusion coefficient of different guest molecules (varying guest lengths and bending stiffness) as a function of tyrosine fractions.

Since the effective size of the stiffer guest molecules increases in comparison to the fully flexible ones, and they do not experience any compaction during the flux, one can hypothesize that this should lead to a slowing down of flux since the guests would now require larger pores to flux through the condensates. To test this hypothesis we measured the time it takes for guests to flux through the host condensates.

Surprisingly, in contrast to our hypothesis, we observed that on average, the stiffer guests (*κ*_B_ = 1.5, 3.0) took less time to flux relative to the fully flexible guests (*κ*_B_ = 0.0) [Fig. 4(b), Fig. S16]. The variations in the flux times were within errors for the shorter guests (G_10_), while the longer guests (G_20_) exhibited notable differences. The small variations for the shorter guests can be attributed to the fact that the pore sizes of the host condensates are comparable to their radius of gyrations. However, for the longer guests, the radius of gyrations are approximately an order larger than the pore sizes which can explain the larger variations of their flux times [Fig. S12]. Additionally, we observed that the slopes of the fitted curves for average flux times decreased with increasing bending rigidities which also indicates that stiffer guests were less sensitive to the decrease in pore sizes of the host condensates.

As in Fig. 2(c), in response to increase in sequence block-iness, we observed a decrease in flux time as a function of sequence blockiness [Fig. S16]. Like before, increasing bending stiffness of the guest molecules decreased the average flux time. However, the difference between the two stiffest values was negligible. We attribute this to the fact that the blocky host sequences lead to structural inhomegeneity of the condensates evidenced through the broader distributions of their pore sizes. The large pores, therefore, allow the guests with varying stiffnesses to flux through indifferently.

To investigate the dynamics in greater detail, we measured the generalized diffusion coefficient of the guest molecules – with varying length and bending stiffness – in both the bulk (equilibrium) and flux (non-equilibrium) simulations [Fig. 4(c)]. In the bulk simulations, varying the bending stiffness of the guest molecules had negligible impact on their diffusivity. However, increase in guest stiffness showed a notable increase in their diffusivity in the flux simulations. This stiffness-induced increase in diffusivity aligns with findings by Zhang et al.,^59^ who demonstrated that thick rods undergo accelerated diffusion within macromolecular networks once the rod length exceeds the mesh size, following sliding dynamics.

Overall, our analysis clearly shows that concentration gradient coupled with molecular bending stiffness leads to an enhanced diffusivity. However, it does not explain the mechanism through which stiffer guests are able to traverse the condensate faster. Therefore, we next performed contact analysis between the host and guest molecules.

### F. Stiffer guests exhibit increased contacts with the host condensates

To characterize intermolecular interactions that potentially affect molecular flux, we performed contact analysis between the host proteins and the fluxing guest molecules. We define two residues to be in contact when they come within 1.2*σ*_*ij*_ of each other, where *i* and *j* refer to the residue types of the interacting pair and *σ*_*ij*_ is their average molecular diameter.

As an example, in Fig. 5(a) we plot the contact map for all the guests with varying bending stiffness for the (Y_2_S_7_)_16_Y_2_S_4_ host condensate (remaining contact maps are in Fig. S17). We observe that regardless of guest length and stiffness, the glycine residues consistently engaged in greater contacts with the tyrosine residues belonging to the host proteins, as reflected by the dark horizontal lines in the contact maps. This is not surprising since the glycine– tyrosine interaction strength is approximately 3 times larger than the glycine–serine interaction. Summing up the contact maps revealed that for all the host proteins that we considered, increase in guest stiffness resulted in a simultaneous increase in the total contacts made by the host and guest molecules [Fig. 5(b,c)]. This implies that stiffer guest molecules experience more interactions with the host proteins as opposed to their flexible counterparts. We believe this apparent increase stems from the fact that the stiffer guests expose a larger surface area which can form contacts with the host proteins, while the flexible guests tend to form collapsed configurations which are even more compact within the host condensate, leading to a smaller exposed surface area.

**FIG. 5:**
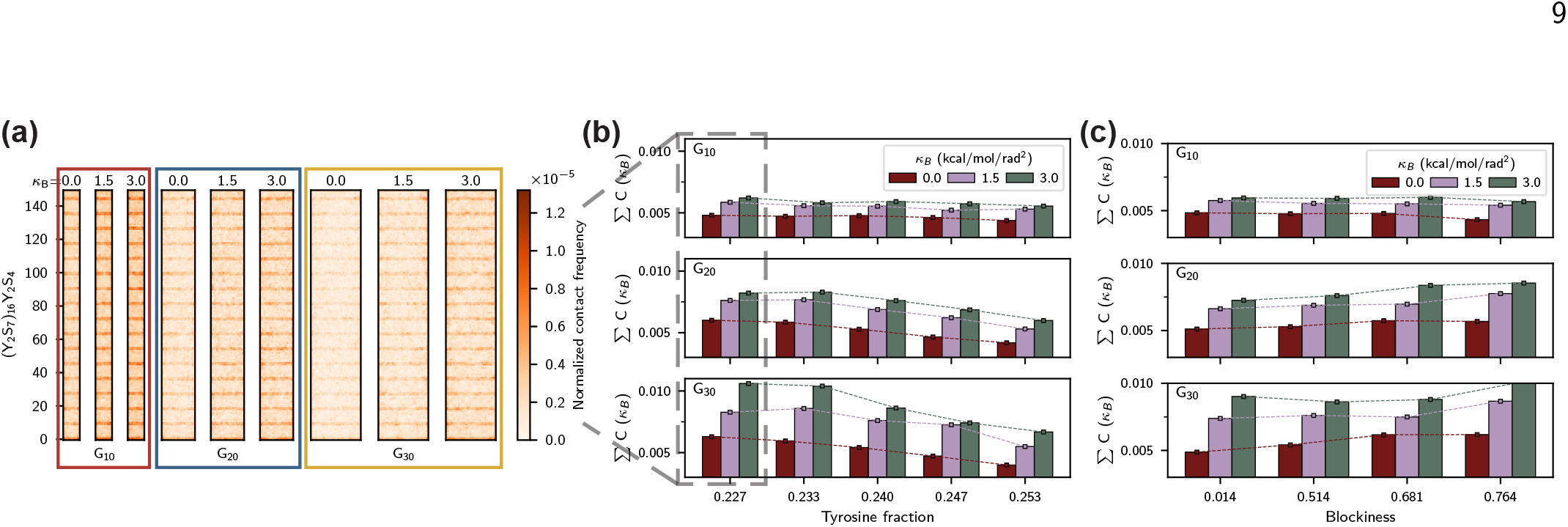
Host-guest contacts increase with guest molecular stiffness. (a) Host-guest contact map for (Y_2_S_7_)_16_Y_2_S_4_ host condensate for all bending rigidities and guest lengths, indicated by colored bounding boxes. (b, c) Total host-guest contact frequencies for all the guest lengths and stiffnesses for host condensates with (b) increasing tyrosine fraction and (c) increasing sequence blockiness.

Interestingly, if we compare the net contacts across the tyrosine fractions, we observed a decline in contacts as the tyrosine fraction increased – pointing to a negative correlation between contact frequency and flux time.

### G. Stronger host–guest interactions hinder molecular flux

To probe the effect of interparticle interactions on molecular flux, we systematically tuned the glycine-tyrosine interaction strength, while keeping all other variables constant. Specifically, the guest molecules were kept fully flexible, and the glycine–tyrosine interaction strength was either (i) reduced, 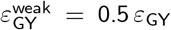, or (ii) increased, 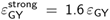. At the lower interaction strength, the glycine–tyrosine interaction is comparable to glycine–serine interaction, whereas at the higher interaction strength, it is equivalent to tyrosine– tyrosine interaction [Fig. 6(a)].

**FIG. 6:**
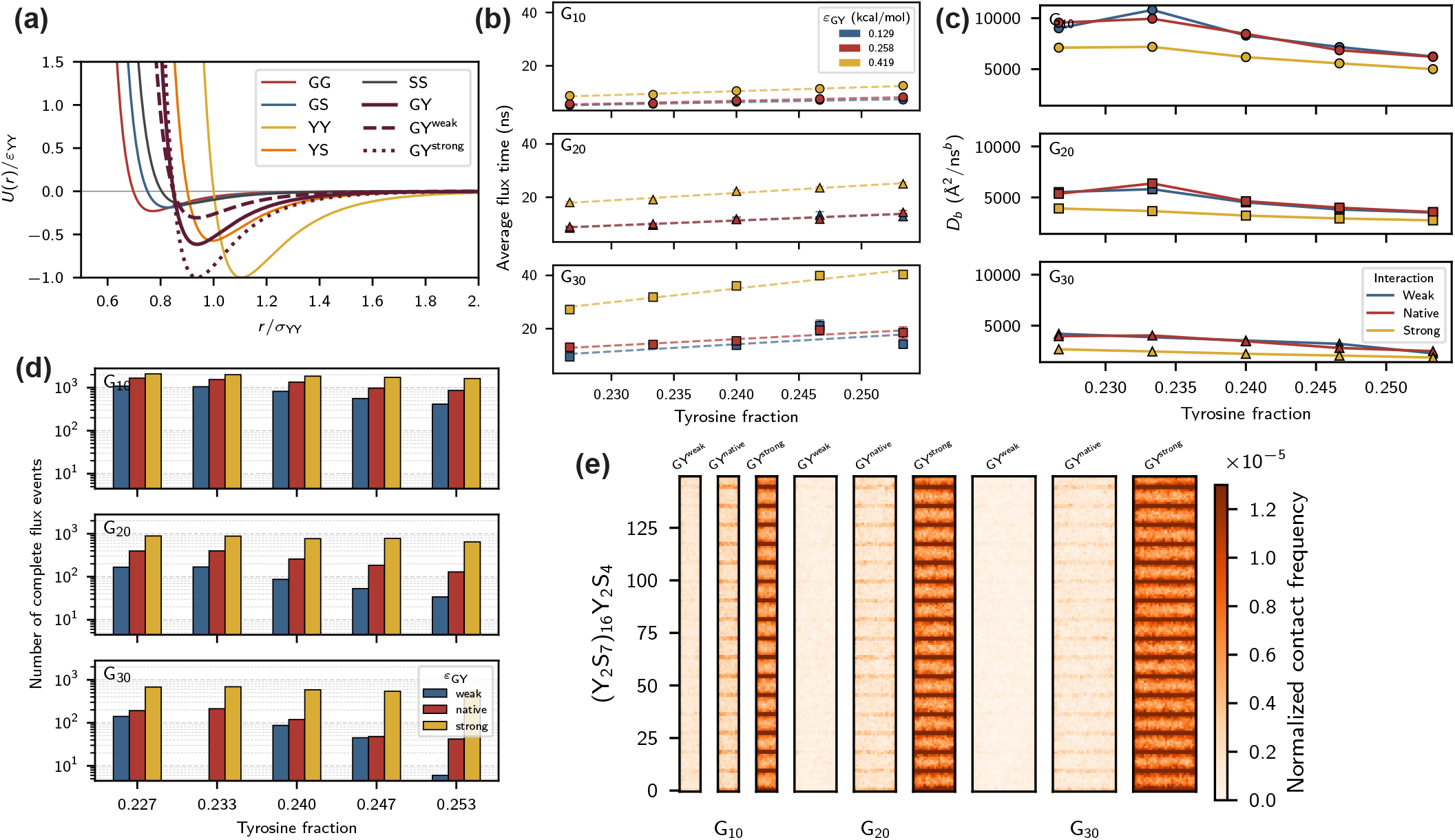
Associative host–guest interactions control diffusion and solubility, therefore determining molecular flux. (a) Pair (Wang-Frenkel) interactions between different residues in the system, with glycine–tyrosine interactions shown in maroon. Solid lines correspond to native Mpipi interactions, while the dashed and dotted curves correspond to reduced and enhanced glycine–tyrosine interactions. (b) Average flux time as a function of tyrosine fraction of the host condensates for G_10_ (top), G_20_ (middle) and G_30_ (bottom) guests. Colors indicate the glycine-tyrosine interaction strengths. (c) Generalized diffusion coefficient as a function of tyrosine fraction for varying interaction strengths. (d) Number of complete flux events as a function of tyrosine fraction for varying interaction strengths. (e) Host-guest contact map for (Y_2_S_7_)_16_Y_2_S_4_ host with varying guests and interaction strengths. Here we note that the color bar maximum is capped at 1.3 × 10^*−*5^ to maintain comparison with contact map from Fig. 5(a). Therefore dark orange may correspond to normalized contact frequencies exceeding the maximum value indicated by the color bar.

We found that stronger host-guest interactions resulted in longer flux times and slower diffusion, whereas the weaker interactions produced results comparable to those observed for the native interactions strength [Fig. 6(b,c)]. Contact analysis further revealed a sharp increase in the host–guest contacts upon strengthening the glycine–tyrosine interaction, accompanied by a decrease in contacts when the interaction strength was reduced [Fig. 6(e)]. Despite the reduction in diffusivity, tracking the total turnover of guest chains that resulted in complete flux revealed that stronger host–guest interactions led to higher throughput, whereas weaker interactions reduced throughput [Fig. 6(d)].

Combining these results with those obtained when the host and guest chains are modified, we can define general principles governing molecular flux through condensates (Table I). First, reducing the tyrosine fraction of the host chains accelerates flux, i.e., decreases the average flux time. As discussed earlier, this follows from the increase in average pore size, which reduces steric hindrance to guest motion. Larger pores also increase guest throughput (the number of complete flux events), as more guests can traverse the condensate. Second, increasing the stiffness of the guest chains also accelerates flux. Stiffer guests adopt more extended conformations with larger surface area, and their rigidity limits conformational entrapment within the host network, allowing them to thread through the pores more efficiently. However, stiffness lowers throughput. This reflects a geometric rather than kinetic constraint: stiffer guests must orient longitudinally (along the long axis of the simulation box) for successful flux events. Notably, both perturbations increase the number of host–guest contacts despite speeding up flux, though for different reasons: larger pores make more of the host chains accessible to the guest, while stiffer guests present more surface area for interaction. Third, rather than modifying the host or the guest, we can tune the cross-interaction strength between them. Strengthening this interaction slows flux and increases contacts, consistent with greater friction between guest and host. The contact number scales with interaction strength, as one would intuitively expect. Throughput increases because stronger cross-interaction leads to higher guest solubility. Taken together, these results demonstrate that molecular flux through condensates is a tunable property governed by several factors.

**TABLE I:** Design principles governing molecular flux through condensates. Each arrow is annotated with the figure that supports it.

| Perturbation | Avg. flux time | Contact freq. | Throughput | Explanation |
| --- | --- | --- | --- | --- |
| ↓ Host tyrosine fraction | ↓ [2(a)] | ↑ [5(b)] | ↑ [6(d)] | Larger pores reduce steric hindrance and expose more of the host chains to the guest. |
| ↑ Guest chain stiffness | ↓ [4(b)] | ↑ [5(b)] | ↓ [S19] | Extended guests present more surface area, while rigidity limits entrapment and lets them thread through pores. |
| ↑ Host–guest interaction strength | ↑ [6(b)] | ↑ [6(e)] | ↑ [6(d)] | Stronger attraction increases effective friction, slowing transit, while more guests engage with the condensate. |

### H. Crosslinking of condensates shrinks pore sizes and therefore hinders molecular flux

Next, we test these design principles against experiments on condensates. Previous work probed molecular transport through the nucleolus by fusing Cry2olig – a domain that oligomerizes under blue light – to the nucleolar protein NPM1, allowing protein crosslinking inside condensates, and measuring dextran mobility by FRAP^50^. The result showed a strong size-dependence: the flux of 70 kDa dextran was minimally affected, whereas transport of 500 kDa dextran was drastically reduced [Fig. 7(a)]. This behavior was attributed to crosslinking reducing the effective pore size of the condensate and thereby hindering diffusion of larger molecules. Using TRACE simulations, we can now test this interpretation directly by asking how crosslinking alters mesh architecture and molecular flux.

**FIG. 7:**
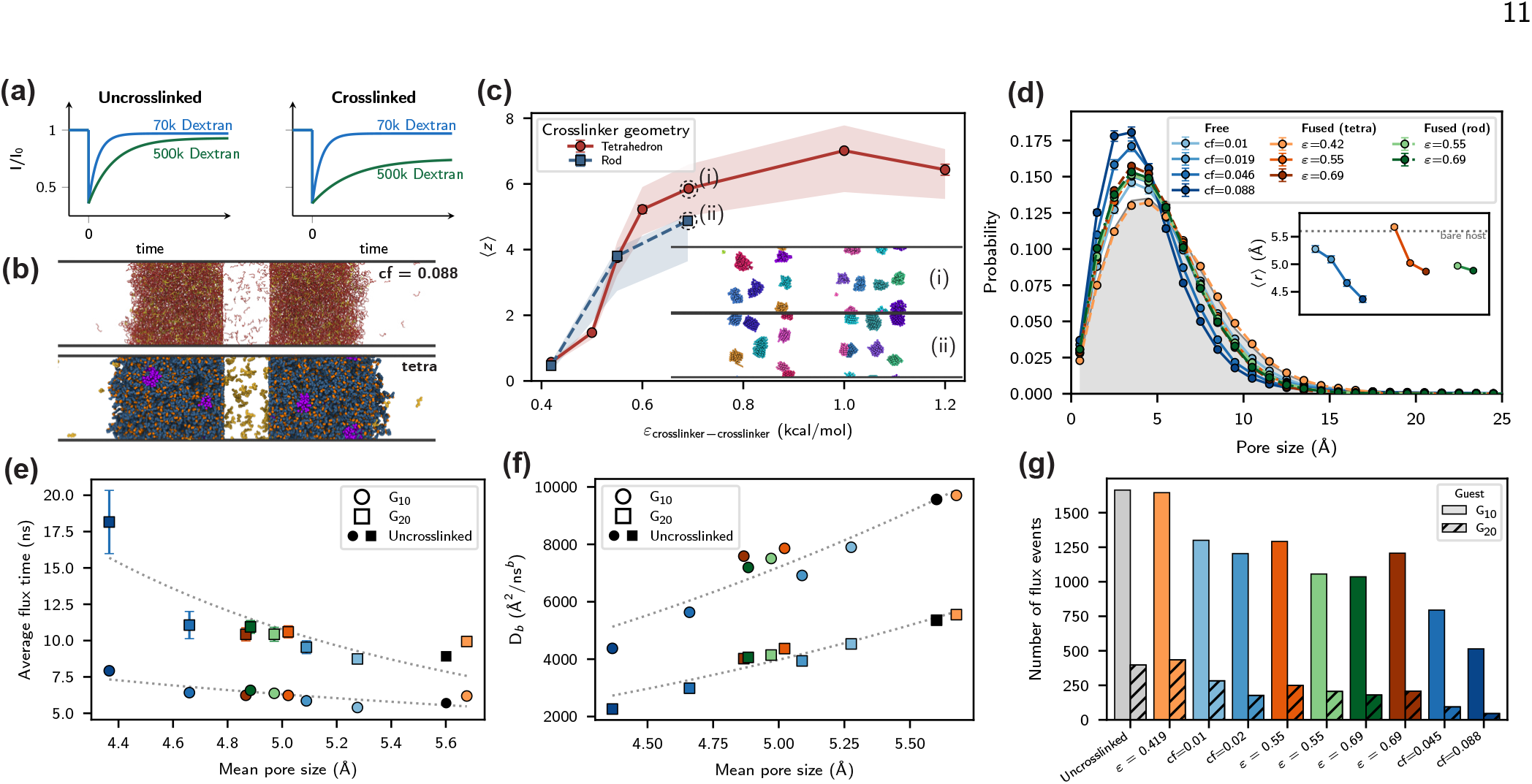
Crosslinking of host condensate leads to shrinkage of pores and therefore hinders flux of guest molecules. (a) Schematic of FRAP recovery curves for dextran chains diffusing through the GC phase of the nucleolus, reproduced from Fig. 5D of Proc. Natl. Acad. Sci. U.S.A. 116 (35).^50^ (b) Simulation snapshots for crosslinked condensates, with top: bonded crosslinking (cf=0.10) and bottom: interaction based crosslinking (tetrahedral). (c) Mean observed valency for the interaction based crosslinkers as a function of interaction strength. Inset snapshots show the clustering of the i. tetrahedral and ii. linear crosslinkers at *ε* = 0.69 kcal/mol. (d) Pore size distribution for the different crosslinking mechanisms – bonded in blues, tetrahedral crosslinkers in oranges, and linear crosslinkers in greens. The original pore size distribution is shown as gray shaded region in the background. Inset shows the mean pore sizes for the different crosslinking mechanisms. (e) Average flux time and (f) generalized diffusion coefficient for glycine guests G_10_ and G_20_ as a function of condensate mean pore size. Dotted lines are guide for eyes. (g) Total flux turnover recorded for the different crosslinked systems in one microsecond simulations. Colors for the scatter points and bars correspond to those in (d).

Rather than model the entire nucleolus – a multicomponent condensate comprising multiple coexisting phases – we design simulations that capture the essential features of the experimental system: the condensate, probe molecules, and a crosslinking mechanism. To that end, we use our model YS proteins as the condensate forming proteins and glycine peptides as the probe molecules. To crosslink the proteins we employed two different methods – (i) crosslinking mediated through ‘free’ crosslinkers, and (ii) crosslinking mediated through crosslinkers ‘fused’ to the condensate-forming proteins.

Following Ref. 60, we insert free crosslinkers into the dilute phase. These crosslinkers partition into the condensate, where each can form up to four irreversible bonds with monomers on distinct protein chains (see Methods). The degree of crosslinking is termed the crosslinking fraction (cf), defined as,

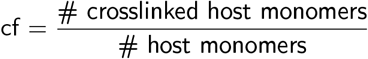

where cf = 1 implies that every particle in the system is crosslinked. In this study we tested different crosslinking fractions (Table III). Fig. 7(b) shows a snapshot of condensate with cf = 0.10.

To more closely recapitulate the experiments, we also introduce crosslinkers fused to the YS proteins. Briefly, the crosslinker interactions are tuned [via *ϵ*, Fig. 7(c)] such that crosslinker–crosslinker interactions are favored over crosslinker–YS interactions, allowing us to mimic domain-mediated oligomerization. Here, we use two geometries for the fused crosslinkers: (i) a centered tetrahedron and (ii) a rod (Fig. S21, Methods). These two geometries allow us to explore the impact of crosslinker topology on the molecular flux. The tetrahedron occupies a bulky volume, whereas the slender rod exposes a larger surface area for interactions. In addition, the rod-like geometry allows for nematic ordering of the crosslinkers.

We quantified crosslinker associations by measuring the mean crosslinker valency which we defined as number of other crosslinkers in contact with any given crosslinker (Methods). We observed an increase in the mean valency with interaction strength, which plateaued around 6, Fig. 7(c), due to excluded volume effects since there are only so many neighbors that one crosslinker can have. We also observed local clustering of crosslinkers at high interaction strengths and nematic ordering of the rods in the clusters, [snapshots in Fig. 7(c)].

We measured the pore size distributions in each crosslinked system [Fig. 7(d)] and, in general, regardless of the crosslinking mechanism we observed a shrinkage in the pore sizes relative to the uncrosslinked condensate (gray shaded region in the plot). Higher crosslinking fraction, in the free crosslinker case, led to smaller mean pore sizes with cf = 0.10 registering a decline in pore sizes by about 22%. The mean pore sizes using fused crosslinkers were in the same range as those obtained from cf=0.02 to cf=0.05 for free crosslinkers. We hypothesized that reduction in the pore sizes should led to reduced flux of the guest molecules.

Indeed we observed an increase in the average flux time with the decrease in the pore sizes, and a concomitant decrease in the generalized diffusion coefficient [Fig. 7(e,f)]. At the highest crosslinking fraction, we observed an almost 50% drop in diffusivity of the guest molecules. We observed decline in the net flux turnover [Fig. 7(g)]; at the highest crosslinking fraction, the net flux dropped by almost 70% and 90% for G_10_ and G_20_ guests, respectively. Importantly, we observe that the transport of the guest molecules showed a strong dependence on the condensate pore size and the different crosslinking mechanisms had negligible impact.

In summary, coupling crosslinking and our TRACE framework allowed us to test the experimental observation that crosslinking hinders the FRAP recovery of large molecules, while leaving the diffusion of small molecules unaffected.

We find that crosslinking alters the pore size of condensates, which in turn affects molecular transport, thereby providing mechanistic insight into the origin of hindered motion in such systems.

## III. DISCUSSION

Material exchange in biomolecular condensates is a fundamental yet poorly understood process, largely due to limitations of existing approaches. In this work, we introduced a novel computational approach (TRACE) to systematically probe molecular flux through condensates with residue-level resolution. By explicitly capturing sequence- and topology-dependent effects, TRACE provides mechanistic insights into how condensate composition and molecular architecture collectively dictate transport properties.

Using condensates composed of tyrosine and serine residues, we isolate and quantify the role of hydrophobic interactions in governing molecular flux behavior. Our results demonstrate that sequence composition and patterning directly shape the effective mesh formed within condensates. Proteins with higher hydrophobic content form denser networks characterized by smaller pore sizes, while blocky sequences form heterogeneous meshes with larger average pores. The pore sizes then dictate molecular flux with smaller pores leading to slower flux.

Analysis of transport behavior revealed that the fluxing molecules exhibit local cage dynamics; for time and length scales smaller than the condensate pores, fluxing molecules moved ballistically, while at longer time scales molecules exhibited diffusive dynamics. This indicates that the fluxing molecules explore their surroundings faster than the condensate pores rearrange, so at short times the pores are effectively static. Once a molecule reaches the edge of its pore and encounter the surrounding host proteins, the dynamics slows down and become diffusive.

Molecule stiffness was found to facilitate their flux through condensates, in line with previous work that reported unconventionally fast dynamics of thick rods through macro-molecular networks.^59^ By modulating host-guest cross interactions, we found that higher friction due to stronger interactions slows down flux but enhance the net throughput. In general, molecular flux depends on the interplay of various factors such as internal organization of the condensates, conformational properties of constituent molecules, and interaction strengths between constituent molecules.

Finally, we leveraged TRACE to probe the effect of crosslinking on molecular flux, finding that crosslinking proteins within condensates leads to shrinking of condensate pores and therefore hindrance to flux. This finding helps illuminate previous experimental findings. In addition, we found that crosslinking mechanism had a minimal impact on flux, which was primarily dependent on condensate pore sizes.

It is important to highlight that the condensates studied here are ‘idealized’. By restricting composition to tyrosine and serine residues, we primarily isolate effects due to hydrophobic forces, but exclude electrostatic interactions that are ubiquitous in condensates and mechanistically distinct in character. Hydrophobic contacts are short-ranged and locally structured, so molecules interact with the network mainly near the pore walls, consistent with the strong dependence of flux on the pore size observed here. Electrostatic interactions are longer-ranged and can create spatially heterogeneous electrostatic environments within a condensate that act as partitioning filters independent of mesh geometry. A charged cargo molecule may be selectively concentrated or excluded from regions of a condensate not because of steric hindrance, but because of charge complementarity or repulsion with the host condensate. Applying TRACE to condensates driven by combination of hydrophobic and electrostatic interactions is, therefore, a natural next step, where charge-based effects may emerge as an orthogonal axis to pore size as physical determinant of molecular flux.

Altogether, TRACE allows us to recast condensate transport as an optimization problem rather than a measurement one. The interplay between different parameters defines a landscape over feature space which can be used to ask how their variations shift the optimum, and in which direction? This opens up an avenue to design condensates with programmable flux behavior for specific cargo with applications in metabolic engineering and therapeutic delivery.

## IV. METHODS

### Mpipi force field

In this model, each amino acid is represented as a single bead centered at the alpha-carbon with a diameter (*σ*), mass, and charge (*q*). The potential energy of the system is defined as,

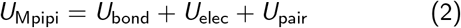

which define the bonded, electrostatic, and short-range non-bonded interactions.

Bonds are described with a harmonic potential,

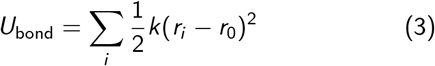

with spring constant *k* = 8.03 Jmol^*−*1^pm^*−*2^ and an equilibrium bond distance *r*_0_ = 381 pm. The charged interactions are described with a screened electrostatic interaction,

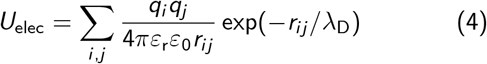

with the dielectric constant of water *ε*_r_ = 80, and Debye screening length *λ*_D_ = 795 pm that corresponds to 0.15 M monovalent salt concentration. For computational efficiency, the electrostatic interaction is cutoff at 3.5 nm.

The pair interaction between any two residues *i* and *j*, separated by distance *r* is described with the Wang–Frenkel (WF) potential,^61^

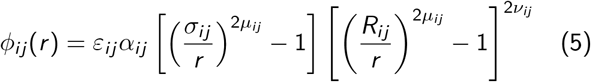

where

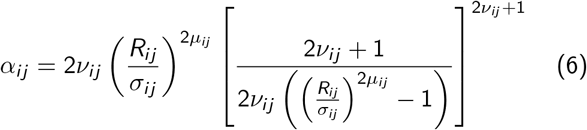

and *ε*_*ij*_ , *σ*_*ij*_ and *µ*_*ij*_ are specified for every unique pair of bead types.^36^ Other parameters are chosen as *ν*_*ij*_ = 1 and *R*_*ij*_ = 3*σ*_*ij*_ . Since the potential goes to zero at *R*_*ij*_ , there is no need to define a cutoff distance for the interaction. The total pair interaction is simply the summation over all particle pairs, *U*_pair_ = Σ_*i* ,*j*_ *ϕ*_*ij*_ .

### Active site

The active site in our system mimics the formation of new molecules and is modeled implicitly. Specifically, we introduce an active site made of discrete particle surfaces that marks the region of material synthesis and is wetted by the surrounding condensate. The discrete particle surfaces allow for the design of active sites with diverse geometries, rendering our framework flexible for use in either slab geometry or with spherical condensates.

The construction of an active site begins with specifying its geometry – planar or spherical in this study. Planar active sites are straightforward to generate by placing particles on a uniformly spaced grid. However, spherical active sites are constructed by recursively subdividing the edges of a regular icosahedron and projecting the vertices onto a sphere, Fig. S23.^58^

The identities of the surface particles are chosen from a probability distribution that reflects the host protein amino acid composition, ensuring wetting of the host condensate due to intermolecular interactions. Within this construct, the interparticle spacing needs to be tuned appropriately, such that the density profile of the host condensate wetting it remains uniform throughout its volume. At small interparticle spacing (4.5, 5.0 Å) the density profiles exhibit a sharp spike close to the active site, indicative of strong attraction of the surface with the host condensate, Fig. S2. While at larger spacings (6.0, 6.5 Å) the density profiles exhibit a dip, indicative of weak attraction of the host condensate to the active site surface, Fig. S2. Ideally, the host condensate density at the active site surface is consistent with the bulk density; here, this occurs when the interparticle spacing was set to 5.5 Å.

A particle-based surface was chosen over a standard Lennard-Jones wall potential as it offered a few key advantages. First, no new interaction parameters are required since the interaction parameters come directly from the force field. Second, with only a single parameter (i.e., interparticle spacing) it produces a density profile matching the bulk dense phase density. It is important that the density of the host condensate in the vicinity of the surface is consistent with the bulk dense phase density to avoid artificial increase in resistance.

### Flux simulations

After constructing the active sites, we place compressed slabs of the host condensates on both sides of the active site, which remains static during the simulations, followed by equilibration for 50 ns. These equilibrated systems serve as reusable configurations to set up initial states for simulations with different guest molecules.

To prepare the initial state for each host–guest combination, multiple copies of the guest molecules were inserted inside the active site and the system was equilibrated for another 30 ns. During the equilibration, guests diffuse out of the active site due to the imposed concentration gradient; diffusing through the host condensate and, in some cases, entering the dilute phase. The equilibrated configuration is then used for subsequent flux simulations.

Production simulations were performed using a hybrid molecular dynamics protocol [Fig. 1(c)], consisting of alternating 20 ns molecular dynamics runs and insertion–deletion steps. In each insertion–deletion step, guest molecules that have entered the dilute phase are reinserted into the active site, thereby maintaining a steady-state concentration gradient. A guest molecule is determined to be in the dilute phase when its center of mass lies outside the condensate interfaces – identified by fitting a super-Gaussian function to the host condensate density profiles. This cycle was repeated 50 times for all the flux simulations, yielding a total simulation time of 1 *µ*s.

All flux simulations were performed in a slab geometry with planar active site surfaces. The simulation box dimensions were 20 × 20 × 200 nm^3^, with the active site surfaces positioned at *z* = 95 nm, and 105 nm.

The number of host protein chains were determined by stipulating that the width of each condensate slab be approximately equal at the simulation temperature of 310 K. Specifically, the condensate thickness was set to 24 nm, slightly larger than the box dimensions along the short axes. The number of host proteins on each side of the active site are,

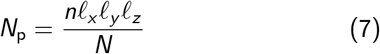

where *n* is the dense phase number density of the condensate at 310 K obtained from direct coexistence simulations, *ℓ*_*x*_ = 20, *ℓ*_*y*_ = 20, *ℓ*_*z*_ = 24 nm are the host condensate slab dimensions, and *N* = 150 is the length of host proteins. Table II lists the number densities and the corresponding number of proteins in the simulations. For the guest molecules, 100, 65, 65 molecules of G_10_, G_20_, G_30_, respectively, were used in all simulations.

**TABLE II:**
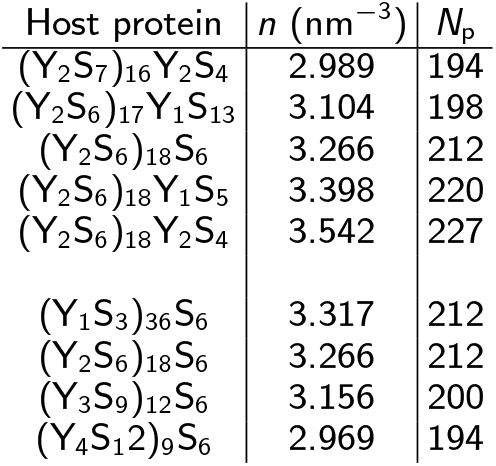
Host condensate dense phase densities and the associated number of protein chains required for a slab of dimensions 20 × 20 × 24 nm^*−*3^.

All simulations were performed in the *NV T* -ensemble at a temperature of 310 K using a Langevin thermostat with a damping factor of 100 ps and a timestep of 10 fs.

### Interparticle interactions

Proper functioning of the flux simulations requires the active site to be semipermeable: host proteins should be excluded from the active site, while guest molecules must be able to permeate it. That is achieved by expanding the number of particle types in the system. Instead of the standard 20 particle types corresponding to the 20 amino acids, we increase the particle types to 60. The first 20 types are used to describe the host molecules, the next 20 represent the active site, and the remaining 20 describe the guest molecules. This separation allows for independent tuning of the interactions between host–guest, host–active site, guest–active site pairs. Here we set the *ε* parameter of the WF potential to zero for guest–active site interactions, thereby making the active site ‘invisible’ to the guest molecules. Fig. S3 shows the exact *ε*_*ij*_ for all the interaction pairs. All other parameters are as given in Ref. 36.

### Isotropic NVT simulations

Isotropic NVT simulations of condensate dense phase with a small number of guest molecules were performed to assess whether compaction of guest molecules arises from their flux dynamics or the host– guest interactions.

A compressed slab of host condensate mixed with a few guest molecules were placed in a cubic simulation box, such that the overall mixture density matched the dense phase density of the host condensate. The host–guest mixture was equilibrated for 40 ns followed by a production simulation for 100 ns where we measured the radius of gyration and mean squared displacement of the guest molecules.

All simulations were performed at 310 K with a simulation box of edge length 21.3 nm. The number of host proteins in the simulation box were the same as in Table II. For the guest molecules, we used 15, 7, and 5 molecules of G_10_, G_20_, G_30_, respectively. These were selected based on the mean number of guest molecules observed within the host condensates during the flux simulations.

### Sequence blockiness

In this study, we define blockiness of the YS sequences as,^56,57^

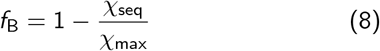

where *χ*_seq_ is the number of Y–S bonds in the sequence and *χ*_max_ = 2 × min(nY, nS) is the maximum number of Y-S bonds possible in a sequence with nY tyrosines and nS serines.

### Average flux time

*t*_flux_, is defined as the time a guest molecule takes to traverse the entire host condensate, entering from the active site and exiting through the opposite side into the dilute phase. For each simulation, we tracked the center-of-mass trajectories of all guest molecules throughout the simulation. For each guest molecule, we identified two key time points: (i) Entry time (*t*_in_): the last recorded frame before the molecule’s center of mass crossed into the dense phase of the host condensate, and (ii) Exit time (*t*_out_): the first recorded frame after the molecule exited the dense phase into the dilute environment.

The flux time for each traversal event was then computed as

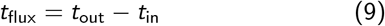

The average flux time was obtained by averaging *t*_flux_ over all traversal events and over all guest molecules in the system. To ensure adequate temporal resolution, molecular positions were sampled every 0.1 ns, which is shorter than the minimum flux times observed across all systems. This sampling frequency ensures that both entry and exit events are resolved with reasonable accuracy.

### Pore size distribution

Polymer theory describes structural properties of polymer solutions using the blob model introduced by de Gennes, in which a blob is a region where interchain interactions are negligible.^62,63^ Within this framework, polymer solutions are categorized into three concentration regimes: dilute solutions, where polymer chains do not overlap; semi-dilute solutions, where chains significantly interpenetrate; and concentrated solutions, where the blob size approaches the polymer bond length.^63,64^

Protein condensates typically correspond to the semi-dilute regime, where the key structural length scale is the effective mesh size. Although prior studies have estimated mesh sizes in biomolecular condensates using monomer density fluctuation correlation lengths, this approach relies on assumptions of homopolymer behavior and structural homogeneity, neither of which hold for biomolecular condensates.^45,65^ To avoid these limitations, we instead quantify mesh sizes using a purely geometrical definition that does not depend on such assumptions.^64,66,67^

Within a geometrical framework, pore volume *V* (*r*) can be defined as the volume of the void space that can be covered by spheres of radius *r* or smaller.^67^ The corresponding pore size distribution can be computed using the following procedure:^64,68^

1. Select a random probe point, **r**_*p*_ in the void.
2. Find the largest sphere of radius *R* such that it encloses **r**_*p*_ and does not overlap with any other particle in the system [Fig. 2(c)]. This can be mathematically formulated as: maximize the function

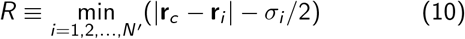

such that,

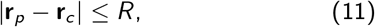

where *σ*_*i*_ is the diameter of *i* ^th^ particle in the system consisting of *N*^*′*^ particles, and **r**_*c*_ is the center of a fitted sphere.
3. Repeat step 1 and 2 many times over multiple condensate configurations (trajectory frames).

The pore size distribution measurements were performed on host condensates in the absence of the guest molecules. Furthermore, to ensure we sampled the interior of the condensate, we defined a buffer distance of 10 Å from the condensate interface.

### Radius of gyration – spatial dependence

To quantify conformational variations of the guest molecules during flux, we measured their radii of gyration. Guest trajectories were saved every 0.1 ns, during which the radius of gyration was computed and binned spatially along the longest axis of the simulation box. The binning was done using the center of mass position of the guest molecules.

The radius of gyration was computed from the gyration tensor, defined as

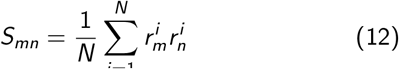

where 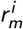 is the m^th^ Cartesian coordinate of the i^th^ particle in a molecule of length *N*. The origin of the coordinate system is chosen to be at the center of mass of the molecule. Diagonalization of the gyration tensor gives the eigenvalues *λ*_1_, *λ*_2_, and *λ*_3_, from which the squared radius of gyration can be computed as

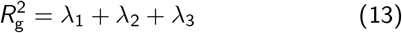

### Mean squared displacement and scaling analysis

To characterize the transport of guests during flux, we measured their displacements while inside the host condensate. To measure dynamics at very short and long time scales, guest molecule trajectories were recorded every 50 fs and 0.1 ns, respectively. Short time simulations were performed for 0.1 ns each with multiple sets of simulations each starting from an unique initial configuration for better statistics. While the long time simulations were performed for 1 microsecond for each host–guest–stiffness combination.

For every guest molecule, time intervals were determined when their center of masses were inside the host condensate. Within each time interval, we computed the MSD of the ceter of mass of the guest molecules, and the MSDs obtained from all guests gathered over an entire 1 microsecond simulation were averaged accordingly.

Transport behavior was classified by fitting the MSD curves to a power law,

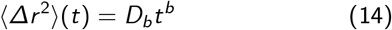

where *D*_*b*_ is the generalized diffusion coefficient and the exponent *b* characterizes the nature of the transport. *b <* 1 indicates sub-diffusive transport, *b* = 1 corresponds to diffusive dynamics, and *b >* 1 indicates super-diffusive behavior.

### Contact analysis

Dominant host–guest interactions were quantified using contact maps. Two particles of types *i* and *j* were considered to be in contact if their separation was less than 1.2*σ*_*ij*_ , where *σ*_*ij*_ refers to the characteristic distance parameter of the Wang–Frenkel potential [Eq. 5].

Contact maps were computed from simulation snapshots taken every 5 ns. The cumulative contact map was normalized by the total number of snapshots *M*, as well as by the number of hosts (*h*) and guests (*g*),

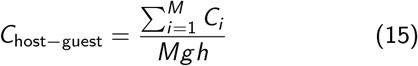

Under this normalization scheme, a value of 1 indicates that a specific pair between host and guest molecules remains in continuous contact throughout the simulation.

### Free crosslinker-based crosslinking

We start from a configuration of an uncrosslinked host condensate that wets the active site, and insert free WCA crosslinkers (glycine sized) in the dilute phase.^69^ After we let the system relax, the free crosslinkers partition into the condensate. Once partitioned, we randomly selected crosslinkers and form irreversible bonds (*k*_cross_ = 8.03 Jmol^*−*1^pm^*−*2^; *r*_0_ = 7 Å) with their nearest monomers within 12 Å such that the bonds are with monomers on unique proteins. Any given crosslinker can have a minimum of two and maximum of four bonds. Having at least two bonds for the crosslinker ensures there is actual crosslinking, while the upper bound of four bonds is an imposed condition.^60^ The number of crosslinkers participating in the crosslinking defines the crosslinking fraction (cf), Table III.

**TABLE III:** Number of crosslinked host monomer and bonds per crosslinking fraction considered in this study. cf is the crosslinking fraction.

| cf | # crosslinked monomers | # crosslink bonds |
| --- | --- | --- |
| 0.01 | 577 | 582 |
| 0.019 | 1142 | 1163 |
| 0.046 | 2716 | 2911 |
| 0.088 | 5137 | 5821 |

### Fused crosslinker-based crosslinking

To achieve crosslinking in experiments, Cry2olig is fused with a target condensate forming protein^50^, (Fig. S20). The Cry2olig oligomerizes when exposed to blue light, which facilitates crosslinking of proteins. Analogously, in the simulations we model fused crosslinkers as small rigid bodies made of five residues that are bonded to the termini of the YS proteins.

We considered two different configurations for the fused crosslinker – (i) centered tetrahedron and (ii) rod [Fig. S21]. The two different geometries allow us to explore the impact of crosslinker geometry on condensate structure and molecular flux. The tetrahedron presents a bulky volume while the rod is slender, which exposes larger surface area for interactions. In addition, the rod-like geometry also allows for nematic ordering of the crosslinkers.

To incorporate a tendency for oligomerisation of the crosslinkers, we carefully tuned their interactions, such that the crosslinkers show preference for other crosslinkers over the YS proteins. To that end, we modeled the crosslinker interactions using the Wang–Frenkel potential with the crosslinker–protein strength *ε*_cp_ = 0.148 kcal*/*mol, equal to the global mean of the interaction strengths in the Mpipi force field (Fig. S22), while scanning a range of values for the crosslinker–crosslinker interaction strength, Fig.7(c).

We quantified the degree of crosslinking by measuring the mean crosslinker valency which we defined as number of other crosslinkers in contact with any given crosslinker. Specifically, two tetrahedra were considered to be in contact when their central monomers were 2.2247*σ* apart, which corresponds to two vertex monomers just touching each other. While for the rods, they were considered to be in contact if any two monomers (belonging to unique rods) were within 1.2*σ*.

## DATA AND CODE AVAILABILITY

Data and code supporting the findings in this study are available at the Joseph group github repository: https://github.com/josephresearch/Biomolecular_flux

## ACKNOWLEDGMENTS

The simulations presented in this article were performed on computational resources managed and supported by Princeton Research Computing, a consortium of groups including the Princeton Institute for Computational Science and Engineering (PICSciE) and Research Computing at Princeton University. The authors thank Dr. Dilimulati Aierken, Pablo Garcia, Nathaniel Hess and Dr. Alina Emelianova for invaluable feedback on the manuscript and figures. The authors also thank other members of the Joseph group for valuable discussions during various stages of the manuscript preparation. Y.M.W. and J.A.J. acknowledge research funding from the Chan Zuckerberg Initiative DAF (an advised fund of Silicon Valley Community Foundation; grant 2023-332391), and the Gordon and Betty Moore Foundation (project no. 10018284).

## AUTHOR CONTRIBUTIONS

Y.M.W.: conceptualization, methodology, investigation, formal analysis, writing – original draft, review and editing, visualization. J.A.J.: conceptualization, methodology, writing – review and editing, supervision, project administration, funding acquisition.

## CONFLICT OF INTEREST

The authors declare no conflict of interests.

## Notes

https://github.com/josephresearch/Biomolecular_flux

### Competing Interest Statement

The authors have declared no competing interest.

### Summary of Updates

Simulations were extended and analyzed to improve statistics. Transport properties analysis was corrected for an error. An additional section was added to showcase the utility of TRACE framework.

## References

[1] S. Alberti and A. A. Hyman, Biomolecular condensates at the nexus of cellular stress, protein aggregation disease and ageing, Nature Reviews Molecular Cell Biology 22, 196 (2021).

[2] A. A. Hyman, C. A. Weber, and F. Jülicher, Liquid-liquid phase separation in biology, Annual Review of Cell and Developmental Biology 30, 39 (2014).

[3] S. Boeynaems, S. Alberti, N. L. Fawzi, T. Mittag, M. Polymenidou, F. Rousseau, J. Schymkowitz, J. Shorter, B. Wolozin, L. Van Den Bosch, et al., Protein phase separation: a new phase in cell biology, Trends in Cell Biology 28, 420 (2018).

[4] C. P. Brangwynne, C. R. Eckmann, D. S. Courson, A. Rybarska, C. Hoege, J. Gharakhani, F. Jülicher, and A. A. Hyman, Germline p granules are liquid droplets that localize by controlled dissolution/condensation, Science 324, 1729 (2009).

[5] S. F. Banani, H. O. Lee, A. A. Hyman, and M. K. Rosen, Biomolecular condensates: organizers of cellular biochemistry, Nature Reviews Molecular Cell biology 18, 285 (2017).

[6] Y. Zhang, A. G. Pyo, R. Kliegman, Y. Jiang, C. P. Brangwynne, H. A. Stone, and N. S. Wingreen, The exchange dynamics of biomolecular condensates, Elife 12, RP91680 (2024).

[7] M. Hondele, S. Heinrich, P. De Los Rios, and K. Weis, Membraneless organelles: phasing out of equilibrium, Emerging Topics in Life Sciences 4, 343 (2020).

[8] J. A. Riback, J. M. Eeftens, D. S. Lee, S. A. Quinodoz, A. Donlic, N. Orlovsky, L. Wiesner, L. Beckers, L. A. Becker, A. R. Strom, et al., Viscoelasticity and advective flow of rna underlies nucleolar form and function, Molecular Cell 83, 3095 (2023).

[9] N. Kedersha, M. R. Cho, W. Li, P. W. Yacono, S. Chen, N. Gilks, D. E. Golan, and P. Anderson, Dynamic shuttling of tia-1 accompanies the recruitment of mrna to mammalian stress granules, The Journal of Cell Biology 151, 1257 (2000).

[10] B. Van Treeck and R. Parker, Principles of stress granules revealed by imaging approaches, Cold Spring Harbor Perspectives in Biology 11, a033068 (2019).

[11] B. Niewidok, M. Igaev, A. Pereira da Graca, A. Strassner, C. Lenzen, C. P. Richter, J. Piehler, R. Kurre, and R. Brandt, Single-molecule imaging reveals dynamic biphasic partition of rna-binding proteins in stress granules, Journal of Cell Biology 217, 1303 (2018).

[12] N. Nag, S. Sasidharan, V. N. Uversky, P. Saudagar, and T. Tripathi, Phase separation of fg-nucleoporins in nuclear pore complexes, Biochimica et Biophysica Acta (BBA)-Molecular Cell Research 1869, 119205 (2022).

[13] M. Stewart, Molecular mechanism of the nuclear protein import cycle, Nature reviews Molecular cell biology 8, 195 (2007).

[14] N. O. Taylor, M.-T. Wei, H. A. Stone, and C. P. Brangwynne, Quantifying dynamics in phase-separated condensates using fluorescence recovery after photobleaching, Biophysical Journal 117, 1285 (2019).

[15] S. Alberti, P. Arosio, R. B. Best, S. Boeynaems, D. Cai, R. Collepardo-Guevara, G. L. Dignon, R. Dimova, S. Elbaum-Garfinkle, N. L. Fawzi, et al., Current practices in the study of biomolecular condensates: a community comment, Nature Communications 16, 7730 (2025).

[16] M. Tokunaga, N. Imamoto, and K. Sakata-Sogawa, Highly inclined thin illumination enables clear single-molecule imaging in cells, Nature Methods 5, 159 (2008).

[17] K. Dörner, M. Gut, D. Overwijn, F. Cao, M. Siketanc, S. Heinrich, N. Beuret, T. Sharpe, K. Lindorff-Larsen, and H. Maria, Tag with caution—how protein tagging influences the formation of condensates, BioRxiv , 2024 (2024).

[18] E. Fatti, S. Khawaja, and K. Weis, The dark side of fluorescent protein tagging—the impact of protein tags on biomolecular condensation, Molecular Biology of the Cell 36, br10 (2025).

[19] S. Chong, T. G. Graham, C. Dugast-Darzacq, G. M. Dailey, X. Darzacq, and R. Tjian, Tuning levels of low-complexity domain interactions to modulate endogenous oncogenic transcription, Molecular Cell 82, 2084 (2022).

[20] G. Gao, E. R. Sumrall, and N. G. Walter, Single molecule tracking reveals nanodomains in biomolecular condensates, BioRxiv , 2024 (2024).

[21] S. Hao, Y. J. Lee, N. B. Goldfajn, E. Flores, J. Liang, H. Fuehrer, J. Demmerle, J. Lippincott-Schwartz, Z. Liu, S. Sukenik, et al., Yap condensates are highly organized hubs, Iscience 27 (2024).

[22] Z. Shen, B. Jia, Y. Xu, J. Wessén, T. Pal, H. S. Chan, S. Du, and M. Zhang, Biological condensates form percolated networks with molecular motion properties distinctly different from dilute solutions, Elife 12, e81907 (2023).

[23] J.-E. Shea, R. B. Best, and J. Mittal, Physics-based computational and theoretical approaches to intrinsically disordered proteins, Current Opinion in Structural Biology 67, 219 (2021).

[24] P. Y. Chew and R. Collepardo-Guevara, Probing molecular and biophysical mechanisms of rna and protein phase transitions with simulations and theory, Current Opinion in Structural Biology 93, 103120 (2025).

[25] M. J. Maristany, A. A. Gonzalez, J. R. Espinosa, J. Huertas, R. Collepardo-Guevara, and J. A. Joseph, Decoding phase separation of prion-like domains through data-driven scaling laws, Elife 13, RP99068 (2025).

[26] N. Galvanetto, M. T. Ivanović, A. Chowdhury, A. Sottini, M. F. Nüesch, D. Nettels, R. B. Best, and B. Schuler, Extreme dynamics in a biomolecular condensate, Nature 619, 876 (2023).

[27] M. J. Maristany, A. Emelianova, P. Y. Chew, A. Aguirre, R. Collepardo-Guevara, and J. A. Joseph, Modeling biomolecular condensates across scales: atomistic, coarsegrained, and data-driven approaches, Advances in physics: X 10, 2592547 (2025).

[28] G. L. Dignon, R. B. Best, and J. Mittal, Biomolecular phase separation: from molecular driving forces to macroscopic properties, Annual review of physical chemistry 71, 53 (2020).

[29] J. Wang, J.-M. Choi, A. S. Holehouse, H. O. Lee, X. Zhang, M. Jahnel, S. Maharana, R. Lemaitre, A. Pozniakovsky, D. Drechsel, et al., A molecular grammar governing the driving forces for phase separation of prion-like rna binding proteins, Cell 174, 688 (2018).

[30] G. L. Dignon, W. Zheng, Y. C. Kim, R. B. Best, and J. Mittal, Sequence determinants of protein phase behavior from a coarse-grained model, PLoS Computational Biology 14, e1005941 (2018).

[31] S. Das, Y.-H. Lin, R. M. Vernon, J. D. Forman-Kay, and H. S. Chan, Comparative roles of charge, π, and hydrophobic interactions in sequence-dependent phase separation of intrinsically disordered proteins, Proceedings of the National Academy of Sciences 117, 28795 (2020).

[32] R. M. Regy, J. Thompson, Y. C. Kim, and J. Mittal, Improved coarse-grained model for studying sequence dependent phase separation of disordered proteins, Protein Science 30, 1371 (2021).

[33] A. R. Tejedor, A. Aguirre Gonzalez, M. J. Maristany, P. Y. Chew, K. Russell, J. Ramirez, J. R. Espinosa, and R. Collepardo-Guevara, Chemically informed coarse-graining of electrostatic forces in charge-rich biomolecular condensates, ACS Central Science 11, 302 (2025).

[34] G. Tesei, T. K. Schulze, R. Crehuet, and K. Lindorff-Larsen, Accurate model of liquid–liquid phase behavior of intrinsically disordered proteins from optimization of single-chain properties, Proceedings of the National Academy of Sciences 118, e2111696118 (2021).

[35] J. M. Lotthammer, G. M. Ginell, D. Griffith, R. J. Emenecker, and A. S. Holehouse, Direct prediction of intrinsically disordered protein conformational properties from sequence, Nature methods 21, 465 (2024).

[36] J. A. Joseph, A. Reinhardt, A. Aguirre, P. Y. Chew, K. O. Russell, J. R. Espinosa, A. Garaizar, and R. Collepardo-Guevara, Physics-driven coarse-grained model for biomolecular phase separation with near-quantitative accuracy, Nature Computational Science 1, 732 (2021).

[37] G. Tesei and K. Lindorff-Larsen, Improved predictions of phase behaviour of intrinsically disordered proteins by tuning the interaction range, Open Research Europe 2, 94 (2023).

[38] P. C. Souza, R. Alessandri, J. Barnoud, S. Thallmair, I. Faustino, F. Grünewald, I. Patmanidis, H. Abdizadeh, B. M. Bruininks, T. A. Wassenaar, et al., Martini 3: a general purpose force field for coarse-grained molecular dynamics, Nature Methods 18, 382 (2021).

[39] M. Boccalini, Y. Berezovska, G. Bussi, M. Paloni, and A. Barducci, Exploring rna destabilization mechanisms in biomolecular condensates through atomistic simulations, Proceedings of the National Academy of Sciences 122, e2425261122 (2025).

[40] N. Galvanetto, M. T. Ivanović, S. A. Del Grosso, A. Chowdhury, A. Sottini, D. Nettels, R. B. Best, and B. Schuler, Material properties of biomolecular condensates emerge from nanoscale dynamics, Proceedings of the National Academy of Sciences 122, e2424135122 (2025).

[41] V. Ramachandran and D. A. Potoyan, Molecular drivers of rna phase separation, Proceedings of the National Academy of Sciences 122, e2511348122 (2025).

[42] P. Y. Chew, J. A. Joseph, R. Collepardo-Guevara, and A. Reinhardt, Thermodynamic origins of two-component multiphase condensates of proteins, Chemical Science 14, 1820 (2023).

[43] A. R. Tejedor, R. Collepardo-Guevara, J. Ramirez, and J. R. Espinosa, Time-dependent material properties of aging biomolecular condensates from different viscoelasticity measurements in molecular dynamics simulations, The Journal of Physical Chemistry B 127, 4441 (2023).

[44] S. Mao, M. S. Chakraverti-Wuerthwein, H. Gaudio, and A. Košmrlj, Designing the morphology of separated phases in multicomponent liquid mixtures, Physical Review Letters 125, 218003 (2020).

[45] D. Sundaravadivelu Devarajan, J. Wang, B. Szała-Mendyk, S. Rekhi, A. Nikoubashman, Y. C. Kim, and J. Mittal, Sequence-dependent material properties of biomolecular condensates and their relation to dilute phase conformations, Nature Communications 15, 1912 (2024).

[46] D. Aierken, S. Aland, S. Bo, S. Boeynaems, D. Cai, S. Carra, L. B. Case, H. S. Chan, J. R. Espinosa, T. K. GrandPre, et al., Roadmap for condensates in cell biology, PRX Life 4, 032501 (2026).

[47] E. Zippo, D. Dormann, T. Speck, and L. S. Stelzl, Molecular simulations of enzymatic phosphorylation of disordered proteins and their condensates, Nature Communications 16, 4649 (2025).

[48] A. Garaizar, J. R. Espinosa, J. A. Joseph, G. Krainer, Y. Shen, T. P. Knowles, and R. Collepardo-Guevara, Aging can transform single-component protein condensates into multiphase architectures, Proceedings of the National Academy of Sciences 119, e2119800119 (2022).

[49] A. Stukowski, Visualization and analysis of atomistic simulation data with OVITO-the Open Visualization Tool, MODELLING AND SIMULATION IN MATERIALS SCIENCE AND ENGINEERING 18, 10.1088/0965-0393/18/1/015012 (2010).

[50] L. Zhu, T. M. Richardson, L. Wacheul, M.-T. Wei, M. Feric, G. Whitney, D. L. Lafontaine, and C. P. Brangwynne, Controlling the material properties and rrna processing function of the nucleolus using light, Proceedings of the National Academy of Sciences 116, 17330 (2019).

[51] S. Sami and S. J. Marrink, Reactive martini: chemical reactions in coarse-grained molecular dynamics simulations, Journal of Chemical Theory and Computation 19, 4040 (2023).

[52] A. Feito, I. Sanchez-Burgos, I. Tejero, E. Sanz, A. Rey, R. Collepardo-Guevara, A. R. Tejedor, and J. R. Espinosa, Benchmarking residue-resolution protein coarse-grained models for simulations of biomolecular condensates, PLOS Computational Biology 21, e1012737 (2025).

[53] D. Nilsson and A. Irbäck, Finite-size shifts in simulated protein droplet phase diagrams, The Journal of Chemical Physics 154 (2021).

[54] G. L. Dignon, W. Zheng, and J. Mittal, Simulation methods for liquid–liquid phase separation of disordered proteins, Current opinion in chemical engineering 23, 92 (2019).

[55] J.-Q. Li, Z.-S. Yan, Y.-Q. Ma, and H.-M. Ding, Sequence-dependent biomolecular phase separation driven by short-range interaction: From material properties to coarsening dynamics, Physical Review E 111, 054403 (2025).

[56] D. Tan, D. Aierken, P. L. Garcia, and J. A. Joseph, Biomolecular condensate microstructure is invariant to sequence-encoded molecular and macroscopic properties, Soft Matter 21, 8635 (2025).

[57] P. L. Garcia and J. A. Joseph, Molecular origins of viscoelasticity in biomolecular condensates, The Journal of chemical physics 164 (2026).

[58] Y. M. Wani, P. G. Kovakas, A. Nikoubashman, and M. P. Howard, Mesoscale simulations of diffusion and sedimentation in shape-anisotropic nanoparticle suspensions, Soft Matter 20, 3942 (2024).

[59] X. Zhang, X. Dai, M. A. Habib, L. Gao, W. Chen, W. Wei, Z. Tang, X. Qi, X. Gong, L. Jiang, et al., Unconventionally fast transport through sliding dynamics of rodlike particles in macromolecular networks, Nature Communications 15, 525 (2024).

[60] T. Yokoyama, Y. Qiang, D. Zwicker, and A. Nikoubashman, Molecular simulations of phase separation in elastic polymer networks, The Journal of Physical Chemistry B 130, 4885 (2026).

[61] X. Wang, S. Ramírez-Hinestrosa, J. Dobnikar, and D. Frenkel, The lennard-jones potential: when (not) to use it, Physical Chemistry Chemical Physics 22, 10624 (2020).

[62] P.-G. De Gennes, Scaling concepts in polymer physics (Cornell university press, 1979).

[63] M. Doi, Introduction to Polymer Physics (Oxford university press, 1996) Chap. Molecular motion in entangled polymer systems.

[64] V. Sorichetti, V. Hugouvieux, and W. Kob, Determining the mesh size of polymer solutions via the pore size distribution, Macromolecules 53, 2568 (2020).

[65] M.-T. Wei, S. Elbaum-Garfinkle, A. S. Holehouse, C. C.-H. Chen, M. Feric, C. B. Arnold, R. D. Priestley, R. V. Pappu, and C. P. Brangwynne, Phase behaviour of disordered proteins underlying low density and high permeability of liquid organelles, Nature Chemistry 9, 1118 (2017).

[66] S. Torquato and M. Avellaneda, Diffusion and reaction in heterogeneous media: Pore size distribution, relaxation times, and mean survival time, The Journal of Chemical Physics 95, 6477 (1991).

[67] L. D. Gelb and K. E. Gubbins, Pore size distributions in porous glasses: a computer simulation study, Langmuir 15, 305 (1999).

[68] S. Bhattacharya and K. E. Gubbins, Fast method for computing pore size distributions of model materials, Langmuir 22, 7726 (2006).

[69] H. C. Andersen, J. D. Weeks, and D. Chandler, Relationship between the hard-sphere fluid and fluids with realistic repulsive forces, Physical Review A 4, 1597 (1971).

