## Supplementary Information for "Non-equilibrium modeling of flux through biomolecular condensates"

### Supplementary Information for
### Non-equilibrium modeling of directed flux through biomolecular condensates

Yashraj M. Wani[1] and Jerelle A. Joseph[1,2,*]

[1]*Department of Chemical and Biological Engineering, Princeton University, Princeton, NJ, USA*

[2]*Omenn–Darling Bioengineering Institute, Princeton University, Princeton, NJ, USA*

(Dated: October 9, 2026)

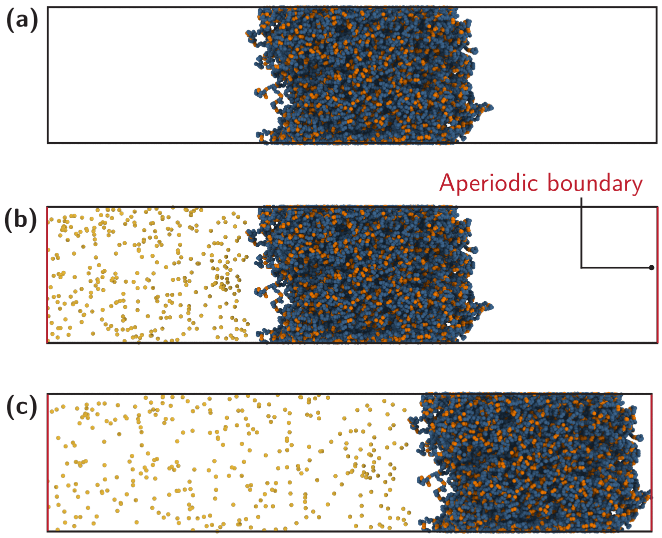


FIG. S1: **Conventional slab simulations can be inadequate for investigating biomolecular flux.** (a) Direct coexistence simulations, commonly employed to examine biomolecular condensates at thermodynamic equilibrium, with periodic boundary conditions along all axes. (b) A possible setup introducing guest molecules (in yellow) on one side of the host condensate to establish a concentration gradient, with aperiodic boundaries along the long axis of the simulation box. (c) In a reorganized configuration, the host condensate drifts to one side of the box, allowing guest molecules to distribute uniformly and reach equilibrium without needing to pass through the condensate. It must be noted here that these snapshots do not represent actual simulations.

Within the slab geometry setup, one can impose a concentration gradient by inserting guest molecules on one side of the condensate, with aperiodic boundaries along the long axis of the simulation box [Fig. S1(b)]. In response, the guest molecules would ideally flux through the condensate toward the dilute region, on the other side. However, this simple configuration suffers from several challenges. First, if the interfacial resistance of the host condensate is large, the system can just reorganize such that the host condensate drifts towards the dilute region, resulting in the depletion of the concentration gradient without the guest molecules traversing the condensate [Fig. S1(c)]. Second, multiphasic condensates in a slab geometry flank the inner phase on either sides and therefore requiring the guest molecules to traverse through the outer phase twice, which can be undesirable. Third, this configuration would constrain us to slab geometries, limiting the flexibility of our approach.

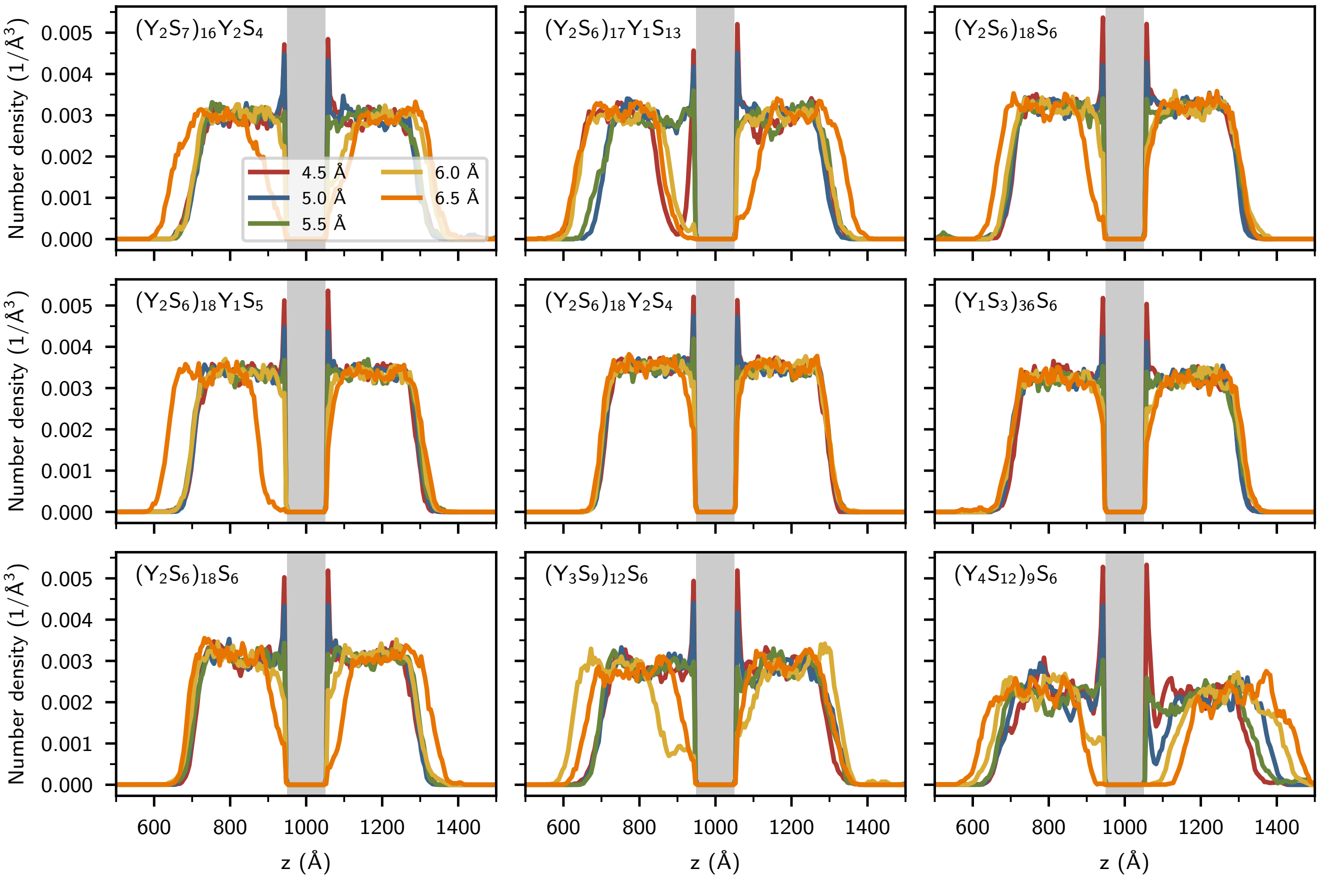


FIG. S2: **Interparticle spacing of the active site surface must be set such that artefacts in the density profiles of the host condensate are prevented.** Number density profiles for all the host condensates with varying interparticle spacing(indicated with different colors) on the active site surface. Gray region highlights the active site.

| Host name | Sequence | Tyrosine fraction |
|---|---|---|
| $(Y_2S_7)_{16}Y_2S_4$ | YYSSSSSSSYYSSSSSSSYYSSSSSSSYYSSSSSSSYYSSSSSSSYY<br>SSSSSSSYYSSSSSSSYYSSSSSSSYYSSSSSSSYYSSSSSSSYYSS<br>SSSSSYYSSSSSSSYYSSSSSSSYYSSSSSSSYYSSSSSSSYYSSSS<br>SSSYYSSSS | 0.2267 |
| $(Y_2S_6)_{17}Y_1S_{13}$ | YYSSSSSSYYSSSSSSYYSSSSSSYYSSSSSSYYSSSSSSYYSSSSS<br>SYYSSSSSSYYSSSSSSYYSSSSSSYYSSSSSSYYSSSSSSYYSSSS<br>SSYYSSSSSSYYSSSSSSYYSSSSSSYYSSSSSSYYSSSSSSYSSSS<br>SSSSSSSSS | 0.2333 |
| $(Y_2S_6)_{18}S_6$ | YYSSSSSSYYSSSSSSYYSSSSSSYYSSSSSSYYSSSSSSYYSSSSS<br>SYYSSSSSSYYSSSSSSYYSSSSSSYYSSSSSSYYSSSSSSYYSSSS<br>SSYYSSSSSSYYSSSSSSYYSSSSSSYYSSSSSSYYSSSSSSYYSSS<br>SSSSSSSSS | 0.2400 |
| $(Y_2S_6)_{18}Y_1S_5$ | YYSSSSSSYYSSSSSSYYSSSSSSYYSSSSSSYYSSSSSSYYSSSSS<br>SYYSSSSSSYYSSSSSSYYSSSSSSYYSSSSSSYYSSSSSSYYSSSS<br>SSYYSSSSSSYYSSSSSSYYSSSSSSYYSSSSSSYYSSSSSSYYSSS<br>SSSYSSSSS | 0.2467 |
| $(Y_2S_6)_{18}Y_2S_4$ | YYSSSSSSYYSSSSSSYYSSSSSSYYSSSSSSYYSSSSSSYYSSSSS<br>SYYSSSSSSYYSSSSSSYYSSSSSSYYSSSSSSYYSSSSSSYYSSSS<br>SSYYSSSSSSYYSSSSSSYYSSSSSSYYSSSSSSYYSSSSSSYYSSS<br>SSSYYSSSS | 0.2533 |

TABLE S1: Host protein sequences with varying Tyrosine fractions.

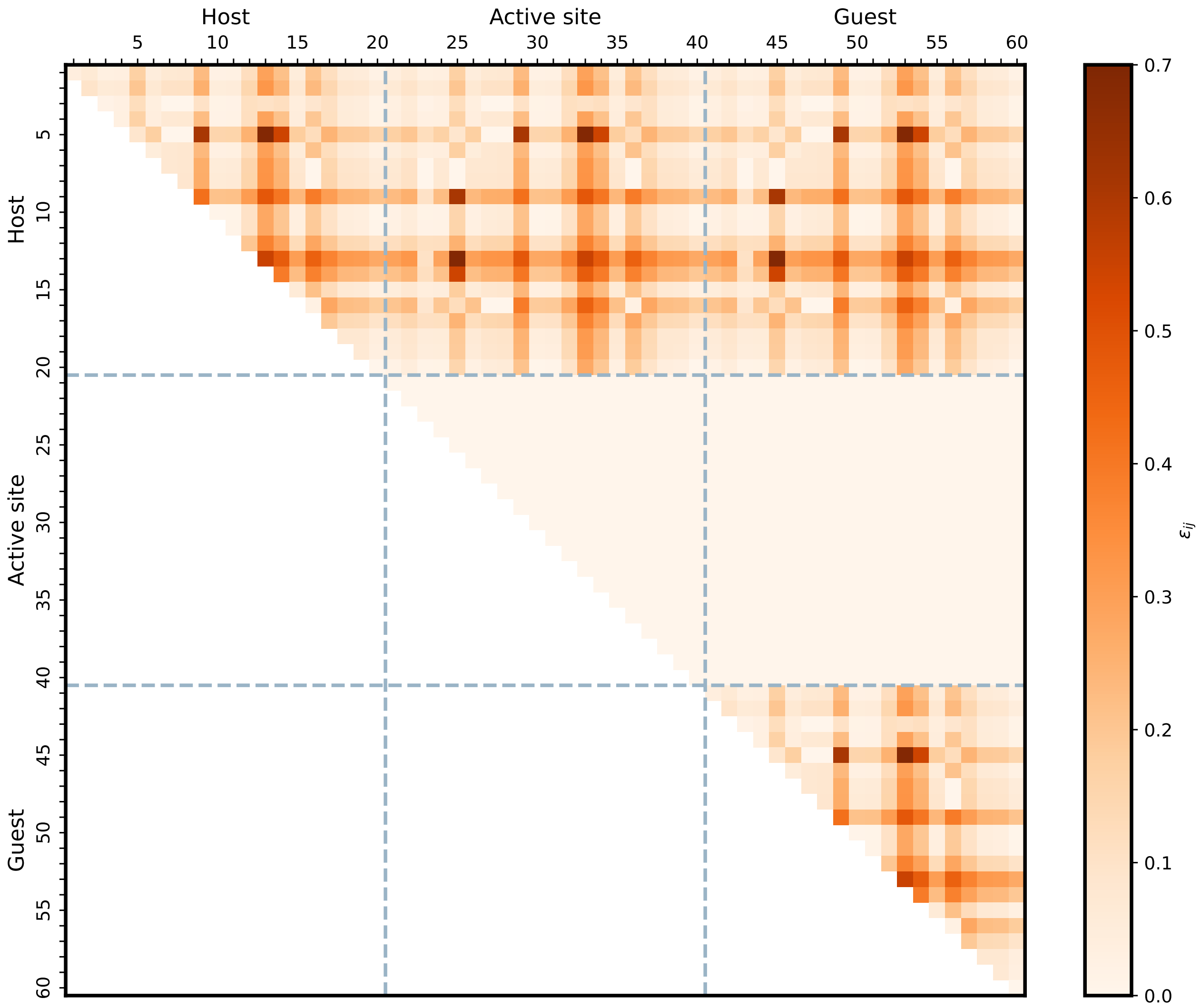


FIG. S3: **Well-depth for the Wang-Frenkel interaction potential.** By setting $\varepsilon_{ij} = 0$ for the interactions between the active site residues and guest residues, the active site surface is rendered 'invisible' to the guest residues.

| Host name | Sequence | Sequence blockiness |
|---|---|---|
| $(Y_1S_3)_{36}S_6$ | YYSSSSSSSYYSSSSSSSYYSSSSSSSYYSSSSSSSYYSSSSSSSYY<br>SSSSSSSYYSSSSSSSYYSSSSSSSYYSSSSSSSYYSSSSSSSYYSS<br>SSSSSYYSSSSSSSYYSSSSSSSYYSSSSSSSYYSSSSSSSYYSSSS<br>SSSYYSSSS | 0.0139 |
| $(Y_2S_6)_{18}S_6$ | YYSSSSSSYYSSSSSSYYSSSSSSYYSSSSSSYYSSSSSSYYSSSSS<br>SYYSSSSSSYYSSSSSSYYSSSSSSYYSSSSSSYYSSSSSSYYSSSS<br>SSYYSSSSSSYYSSSSSSYYSSSSSSYYSSSSSSYYSSSSSSYSSSS<br>SSSSSSSSS | 0.5139 |
| $(Y_3S_9)_{12}S_6$ | YYSSSSSSYYSSSSSSYYSSSSSSYYSSSSSSYYSSSSSSYYSSSSS<br>SYYSSSSSSYYSSSSSSYYSSSSSSYYSSSSSSYYSSSSSSYYSSSS<br>SSYYSSSSSSYYSSSSSSYYSSSSSSYYSSSSSSYYSSSSSSYYSSS<br>SSSSSSSSS | 0.6805 |
| $(Y_4S_12)_9S_6$ | YYSSSSSSYYSSSSSSYYSSSSSSYYSSSSSSYYSSSSSSYYSSSSS<br>SYYSSSSSSYYSSSSSSYYSSSSSSYYSSSSSSYYSSSSSSYYSSSS<br>SSYYSSSSSSYYSSSSSSYYSSSSSSYYSSSSSSYYSSSSSSYYSSS<br>SSSYSSSSS | 0.7639 |

TABLE S2: Host protein sequences with varying sequence blockiness.

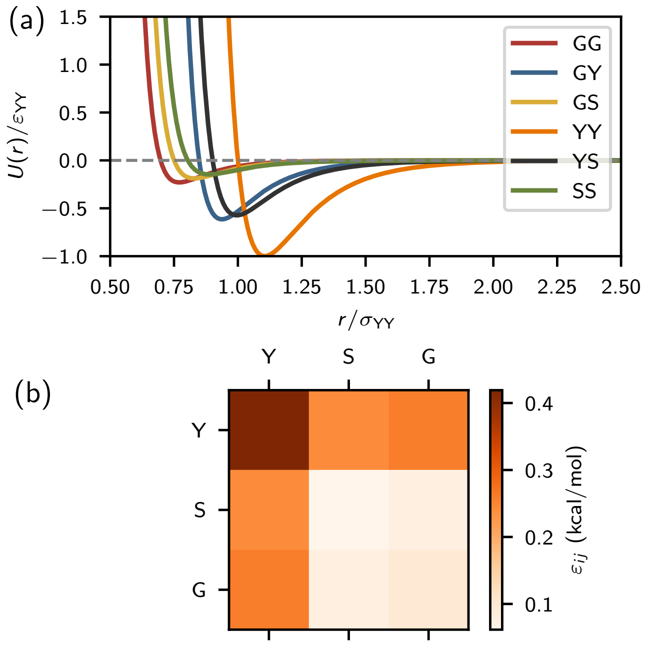


FIG. S4: **Effective three-site interaction model.** (a) Wang-Frenkel interaction potential energy normalized by the Y-Y interaction strength $\varepsilon_{YY}$ and $\sigma_{YY}$. (b) Interaction strength matrix showing the absolute well-depths for the different interaction pairs.

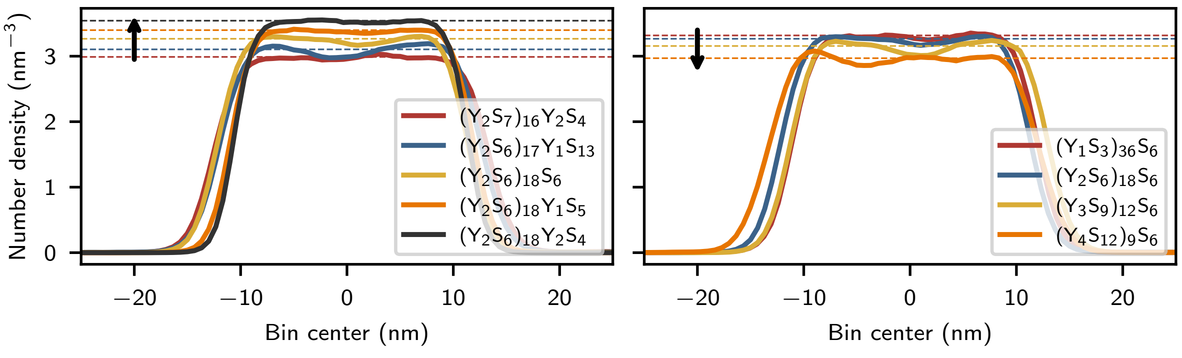


FIG. S5: **Increase in tyrosine fraction leads to increase in dense phase density, while increase in chain blockiness leads to decrease in the dense phase densities.** Number density profiles from direct-coexistence simulations for the host sequences at 310 K. Left: Increasing tyrosine fraction; Right: Increase in blockiness.

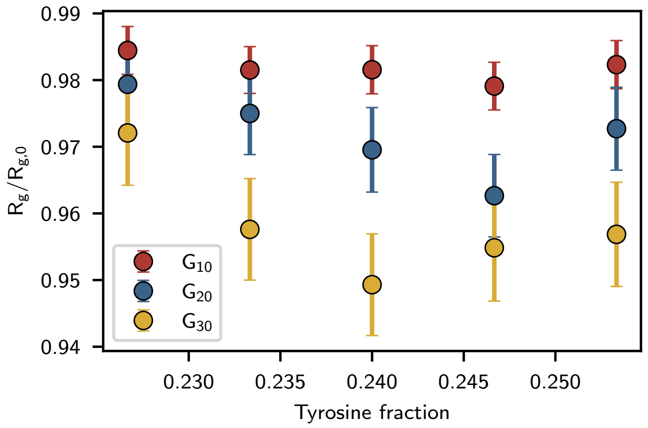


FIG. S6: **Host condensate environment acts as poor solvent for glycine peptides.** Normalized radius of gyration of guest peptides of varying lengths (10, 20, and 30 residues) within the dense phase formed by host sequences with different tyrosine fractions. Radius of gyration values are normalized relative to those measured in the dilute phase.

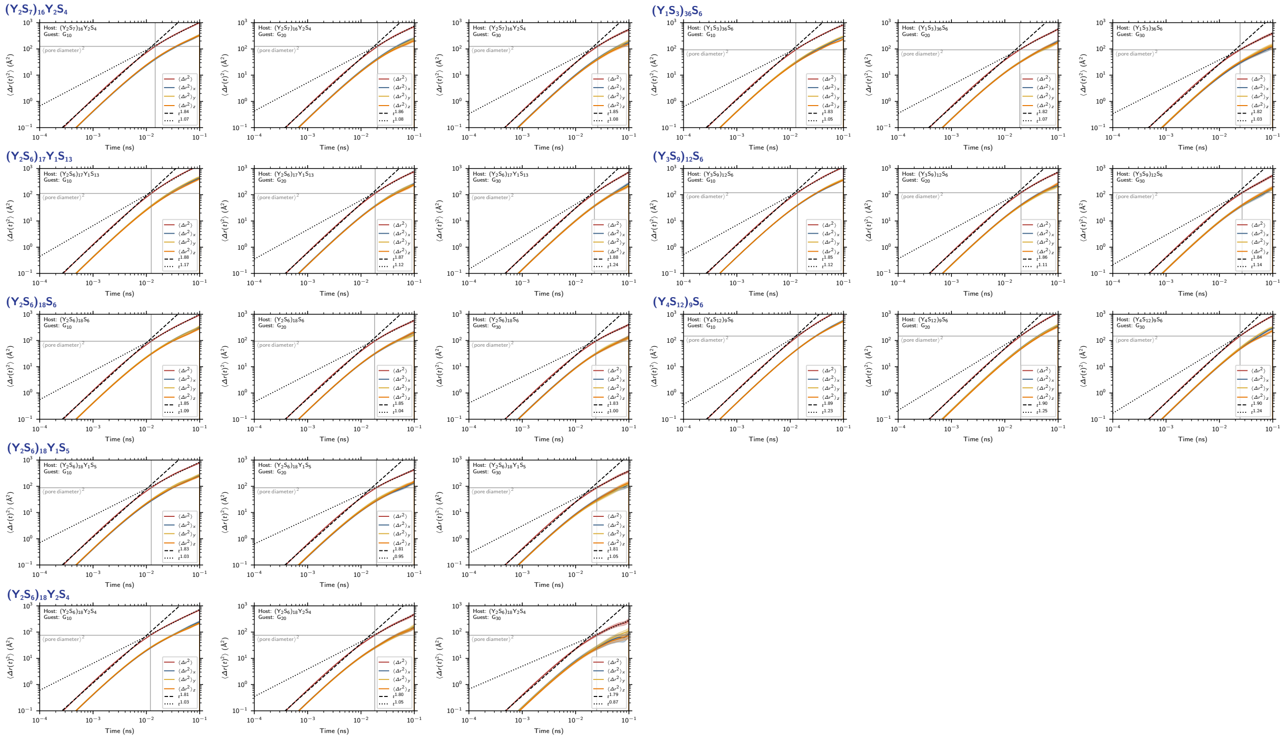


FIG. S7: **Guest molecules exhibit crossover from ballistic to diffusive dynamics at length scale corresponding to the pore diameters of the host condensates.** Mean squared displacement for all host guest combinations at $t < 0.1$ ns. Dashed and dotted lines correspond to ballistic regime and diffusive regimes, respectively.

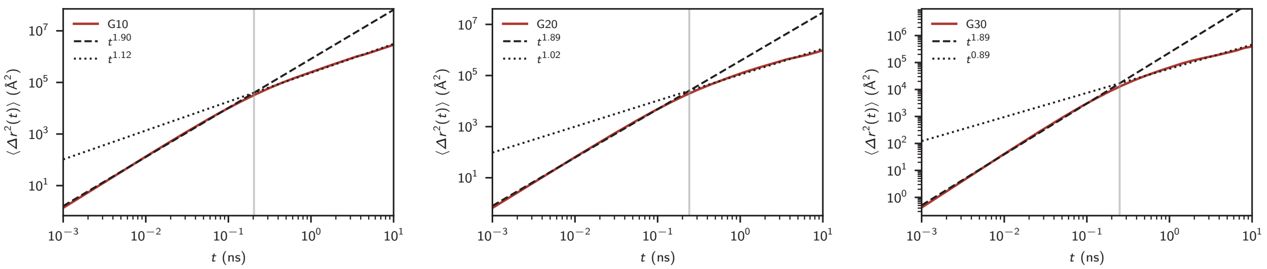


FIG. S8: **The intrinsic ballistic to diffusive crossover stemming from Langevin equations of motion, occurs at $t > 0.1$ ns for the guest molecules, validating the local cage dynamics observations in Fig. S7.** Mean squared displacement for guests at $t < 10$ ns. Dashed and dotted lines correspond to ballistic regime and diffusive regimes, respectively.

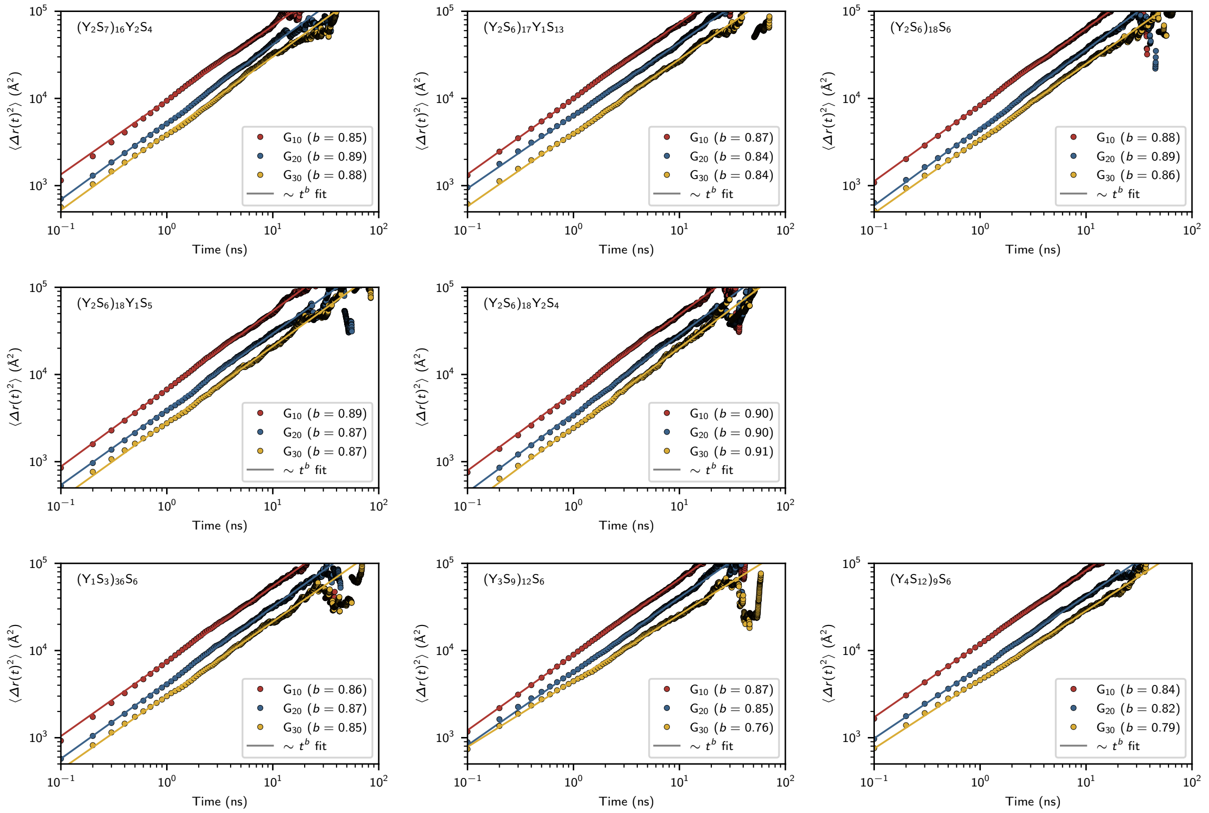


FIG. S9: **Guest molecules exhibit diffusive dynamics at times $t > 0.1$ ns.** Center of mass mean squared displacement for all host guest combinations at $t > 0.1$ ns. Solid lines are guide for eye indicating $\sim t^b$ scaling.

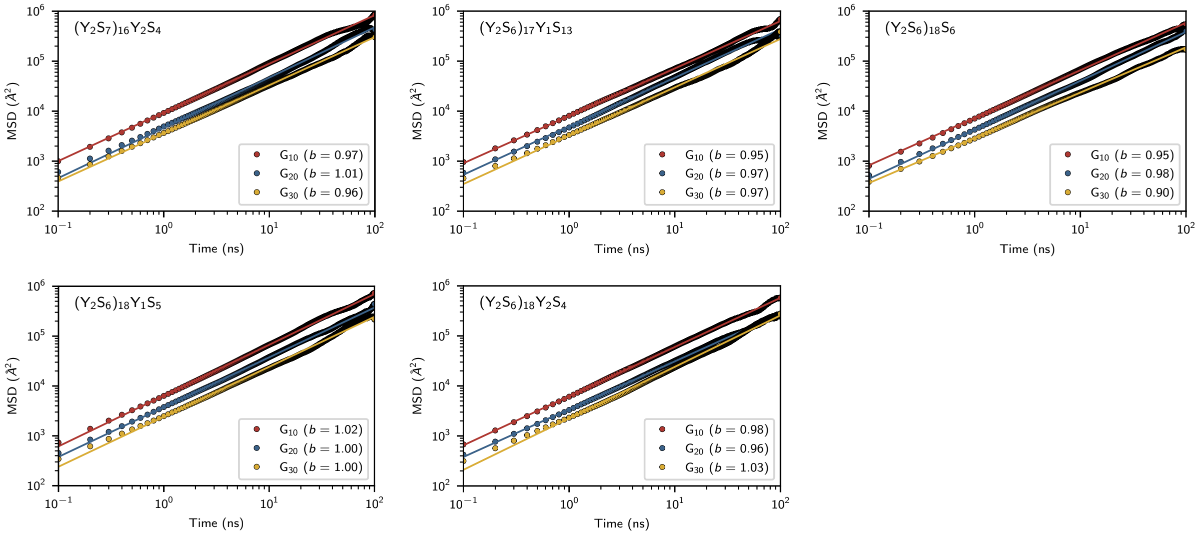


FIG. S10: **Mean squared displacement of guest molecules in NVT simulations.** Solid lines are guide for eye indicating $\sim t^b$ scaling.

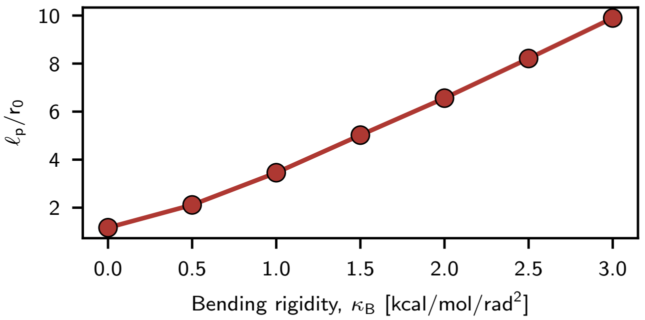


FIG. S11: **Persistence length, $\ell_p$, of glycine peptides as a function of bending stiffness,** $\kappa_{\mathrm{B}}$, normalized w.r.t. the equilibrium bond length, $r_0$. At the highest bending stiffness glycine peptides $N_{\mathrm{m}} = 11$ behave as rods.

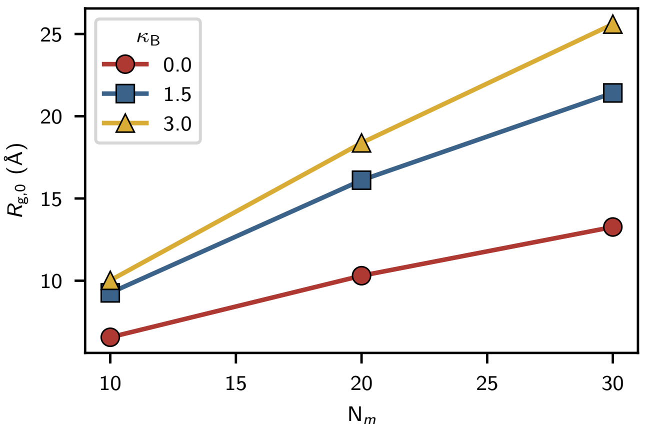


FIG. S12: **Single chain radius of gyration of glycine peptides with varying lengths (x-axis) and bending rigidities (different colors).**

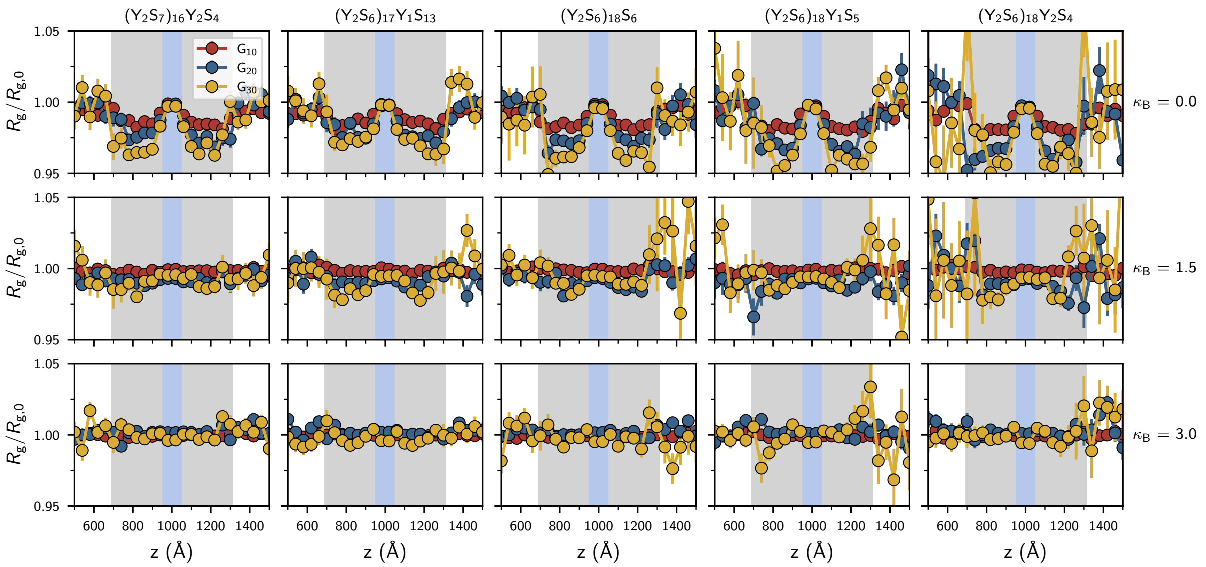


FIG. S13: **Stiffer guest molecules experience lower compaction relative to their dilute phase radius of gyration, in host condensates with increasing tyrosine fraction.** Spatial dependence of radius of gyration of guest peptides as they flux through host condensates with varying tyrosine fraction (increasing from left to right) with increasing bending stiffness of the guest peptides (top to bottom). $R_g$ values are normalized with their respective dilute/single chain values.

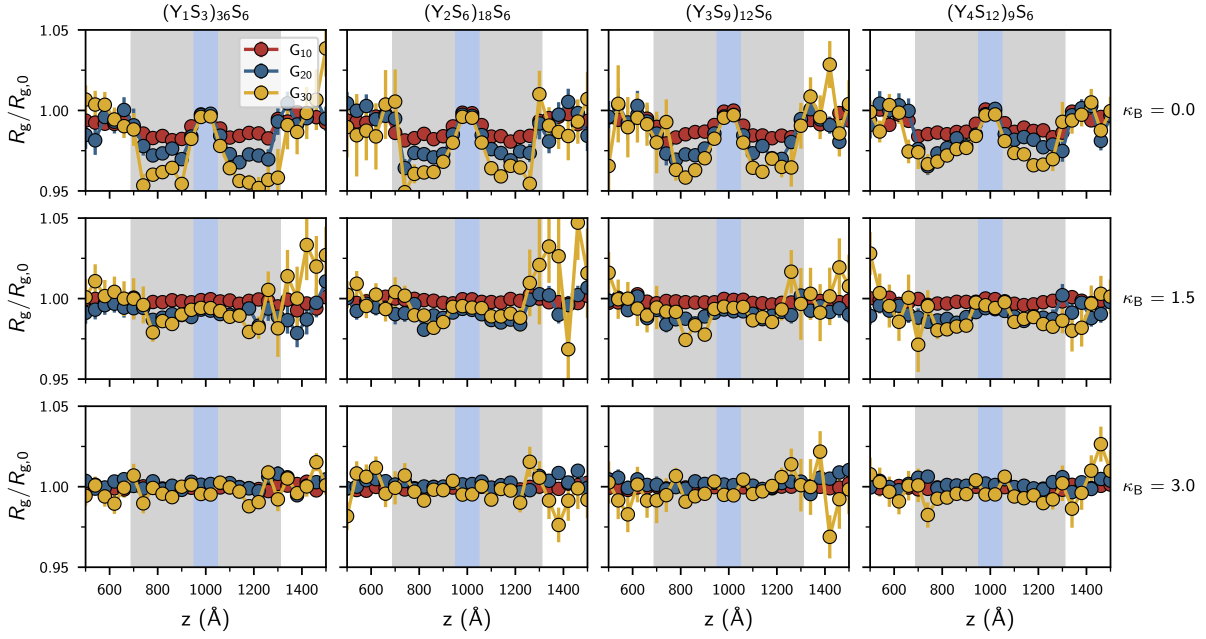


FIG. S14: **Stiffer guest molecules experience lower compaction relative to their dilute phase radius of gyration, in host condensates with increasing blockiness.** Spatial dependence of radius of gyration of guest peptides as they flux through host condensates with varying sequence blockiness (increasing from left to right) with increasing bending stiffness of the guest peptides (top to bottom). $R_g$ values are normalized with their respective dilute/single chain values.

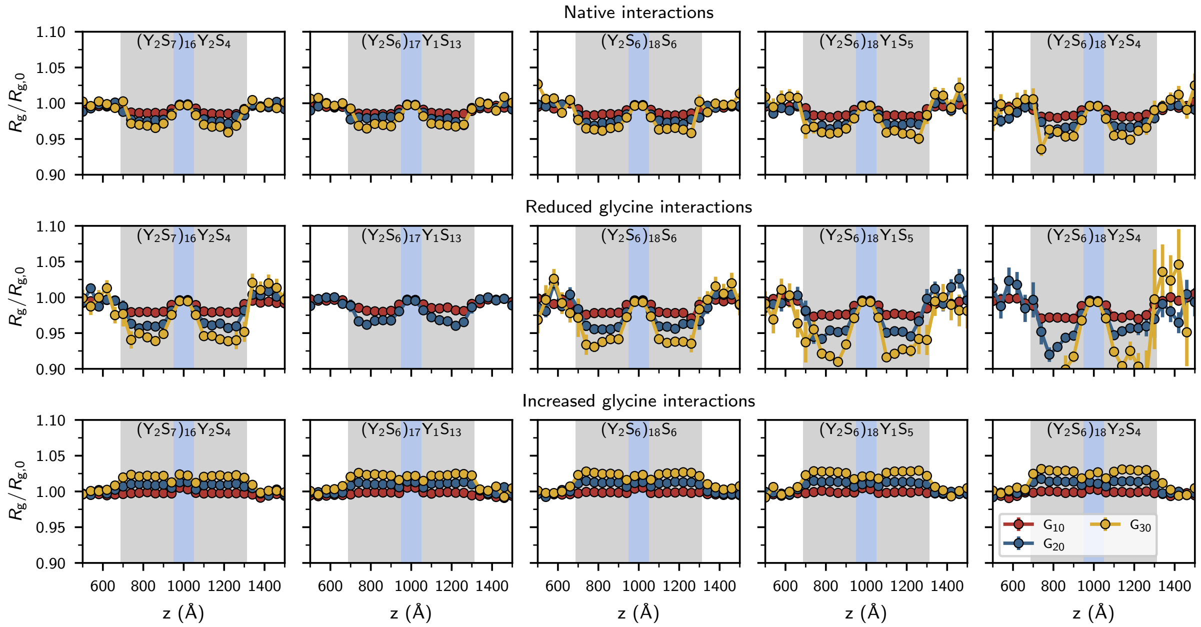


FIG. S15: **Guest molecule conformations are dictated by the solvent quality that the host condensate presents.** Spatial dependence of radius of gyration of guest peptides as they flux through host condensates with varying Glycine–Tyrosine interaction strengths: Native Mpipi interaction strength (Top), $R_g$ values are normalized with their respective dilute/single chain values.

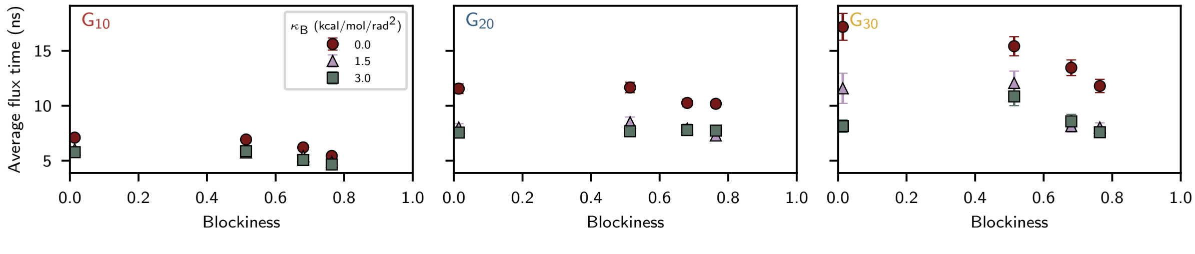


FIG. S16: **Average flux time as a function of host sequence blockiness**, with different guest molecules – left: $G_{10}$, middle: $G_{20}$, and right: $G_{30}$. Different colors correspond to varying guest stiffness.

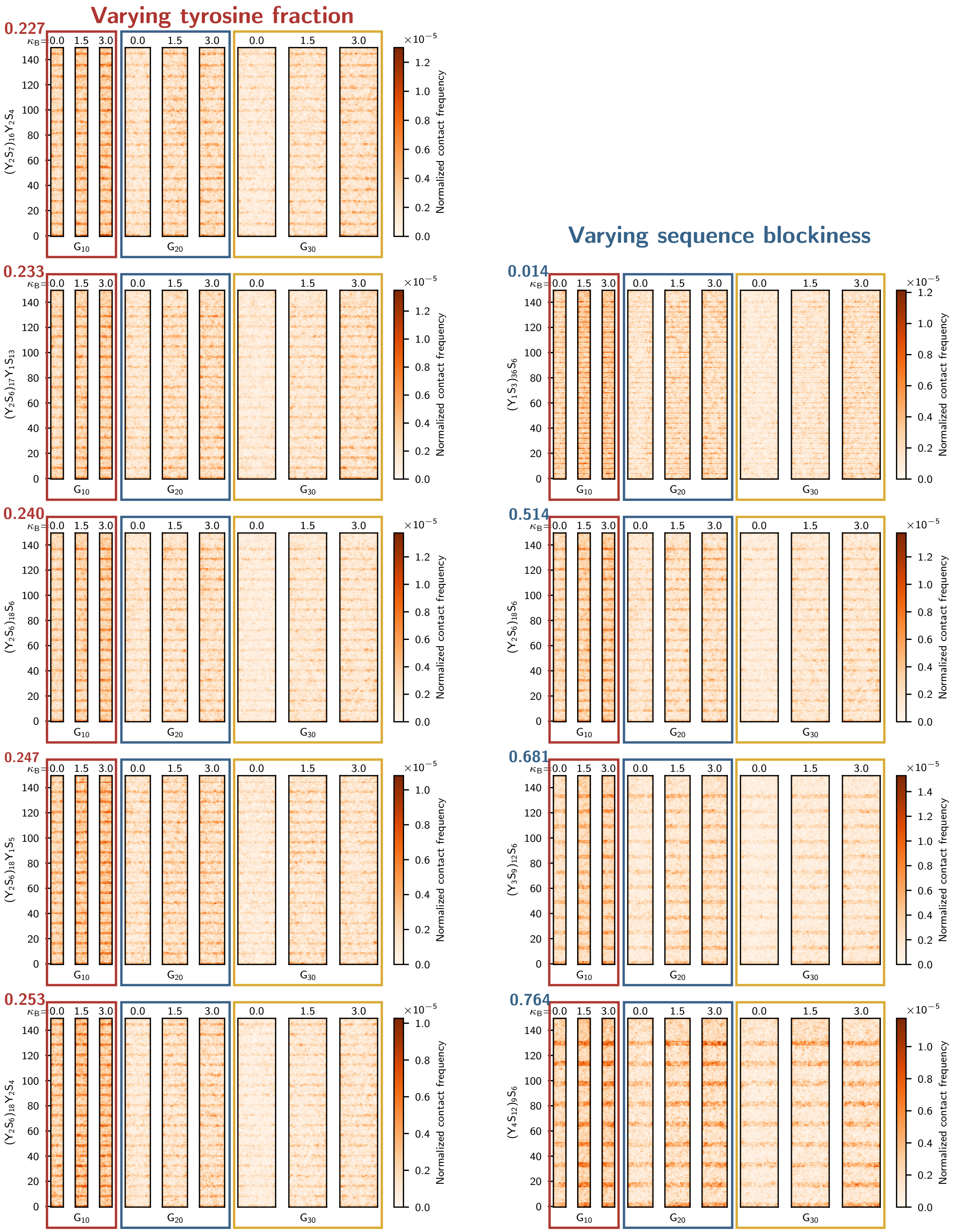


FIG. S17: **Host-guest contact maps show higher contacts between glycine and tyrosine residues, with the contact frequencies increasing with increasing guest stiffness.** Host-guest contact maps for all the host sequences investigated with varying tyrosine fraction (left) and varying sequence blockiness.

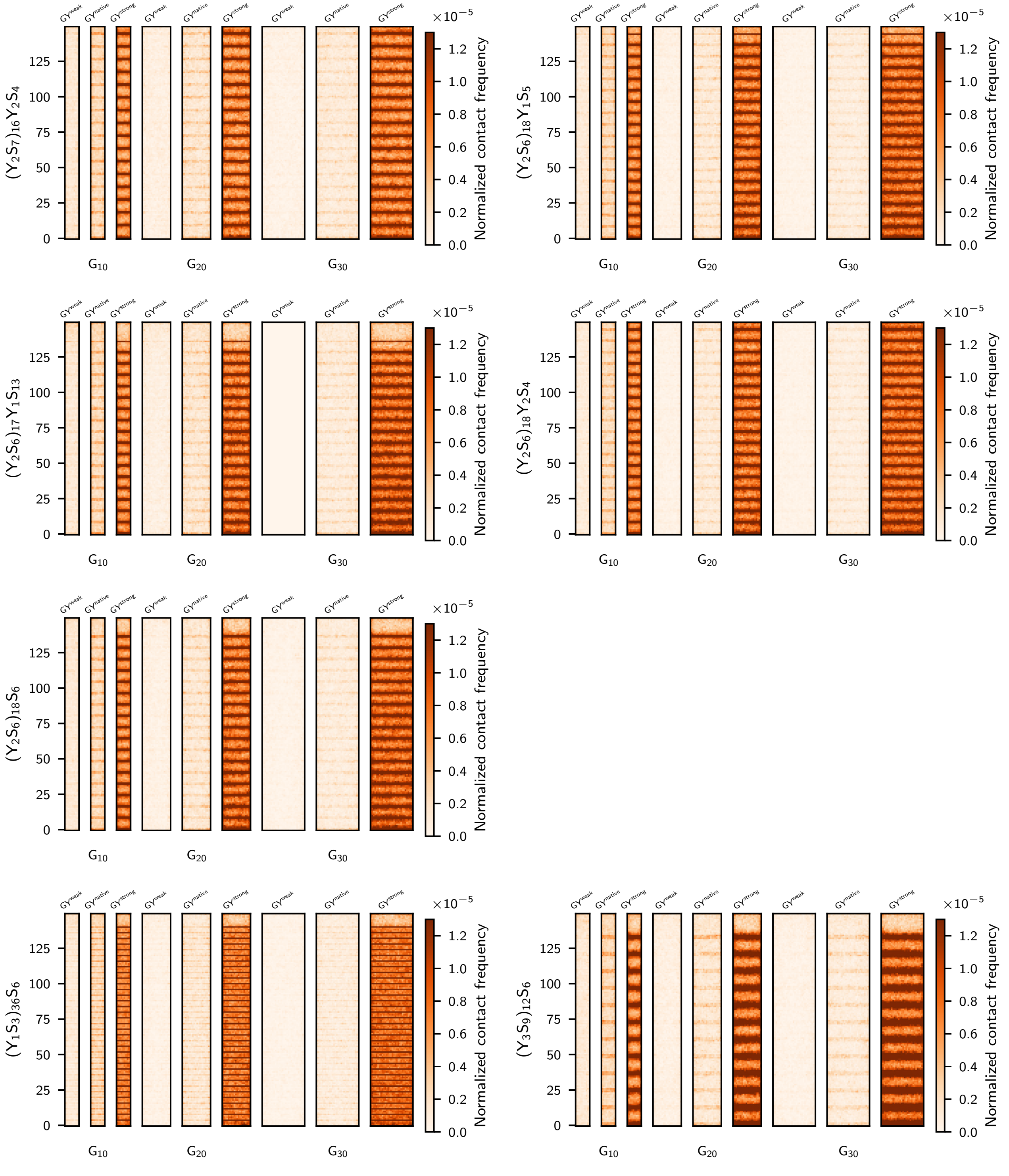


FIG. S18: **Associative interactions lead to increased host-guest contacts.** Contact maps for different host condensates with varying guests and glycine–tyrosine interaction strengths.

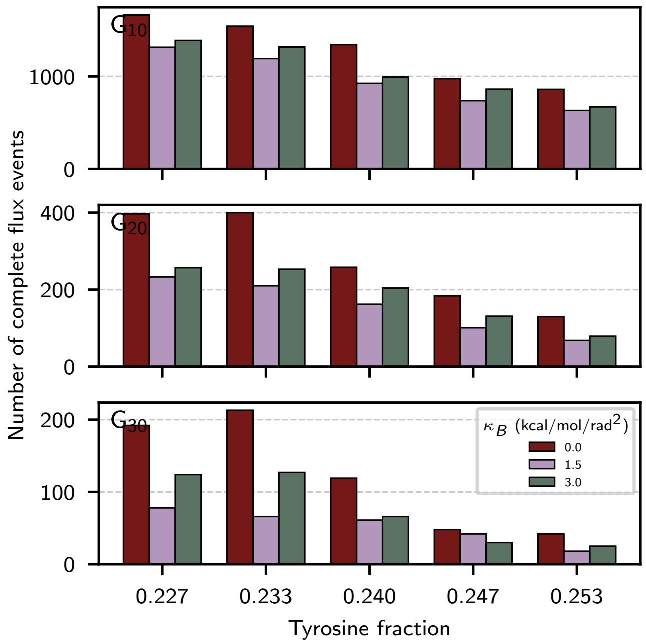


FIG. S19: **Increasing guest stiffness reduces flux turnover.**

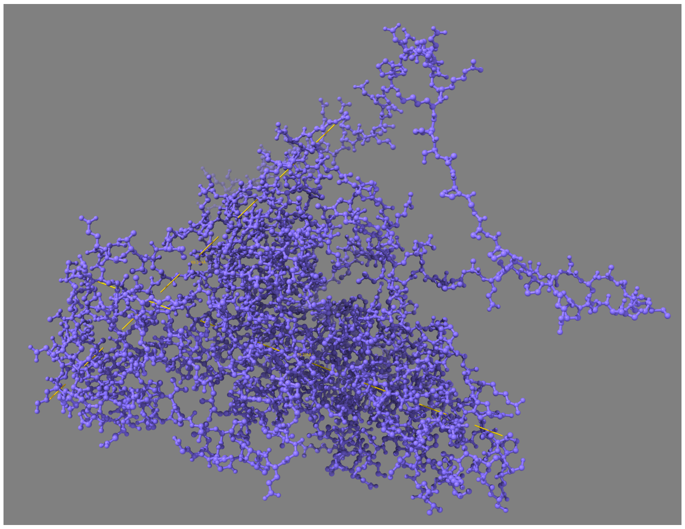

FIG. S20: **Cry2PHR domain of Cry2 protein extends approximately 7 nm.** Uniprot: Q49AN0-2

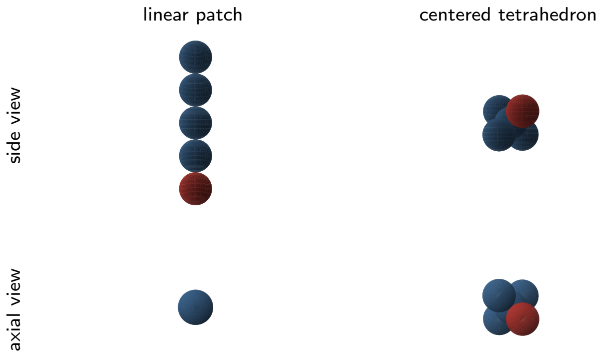


FIG. S21: **Explicit crosslinker geometries – (i) rod-like geometry and (ii) centered tetrahedron.**

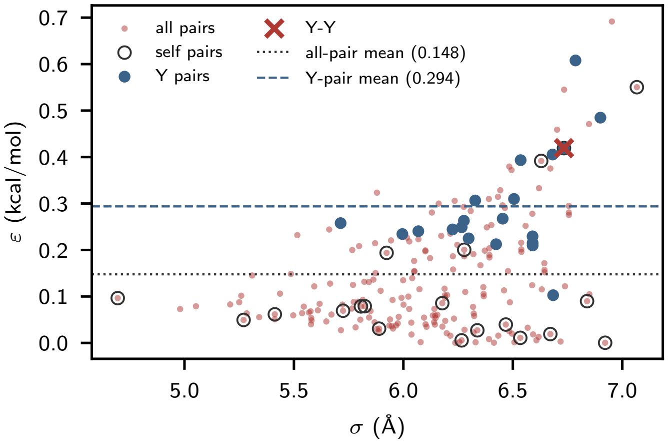


FIG. S22: **Mpipi force-field landscape plotting $\sigma$ and $\varepsilon$ in the Wang-Frenkel interaction for all the interaction pairs**

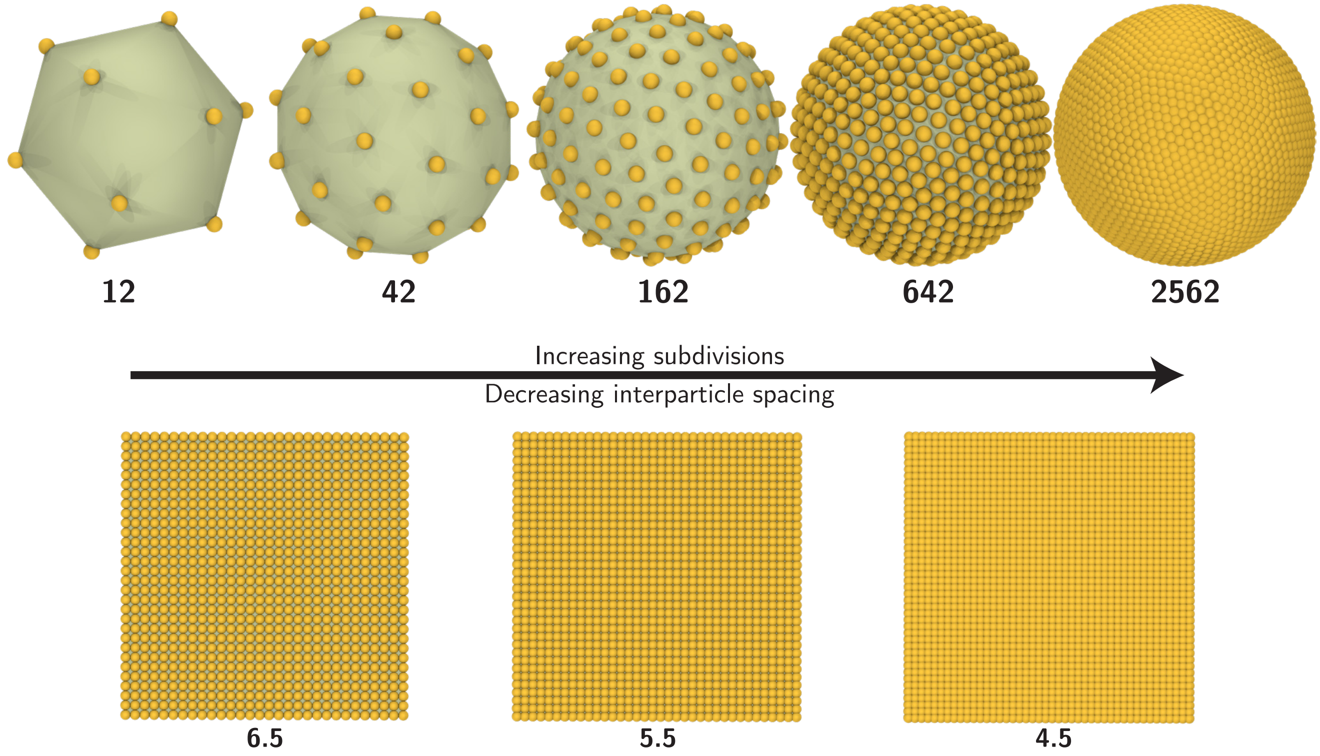


FIG. S23: **Discrete particle models for spherical and planar active site surfaces.** Spherical and planar active sites defined using discrete particle model with increasing level of subdivisions and decreasing interparticle spacings, respectively. Spherical active site: starting from an icosahedron (subdivisions = 0) we have an active site consisting of 12 vertices and 30 edges. With every subdivision, the number of edges quadruple.

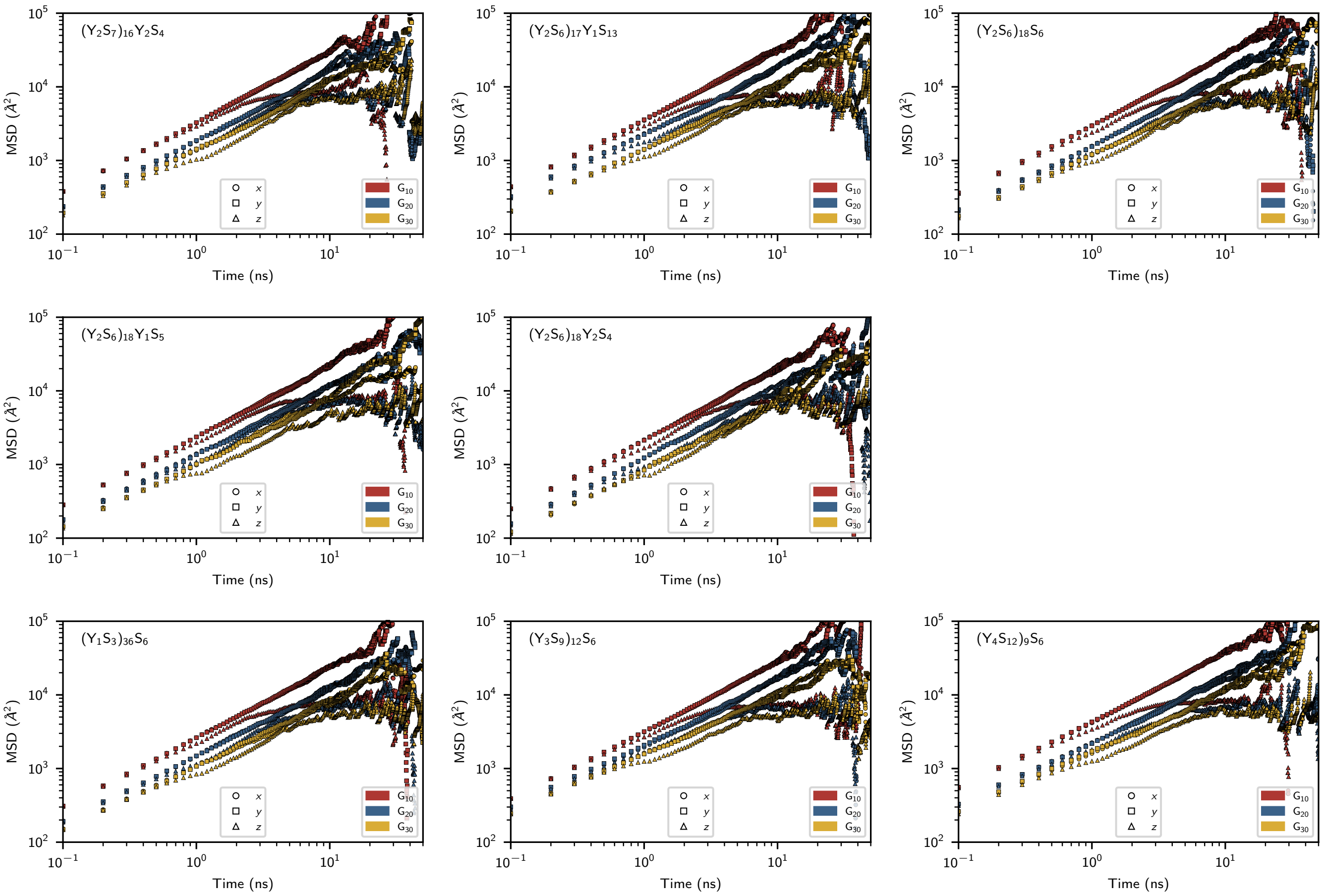


FIG. S24: **Mean squared displacement of guest molecules decomposed along the x, y, z directions.** The MSD deviates from the x and y direction since the slab is finite along the z-direction while along the x and y directions the periodic boundaries wrap the condensates.

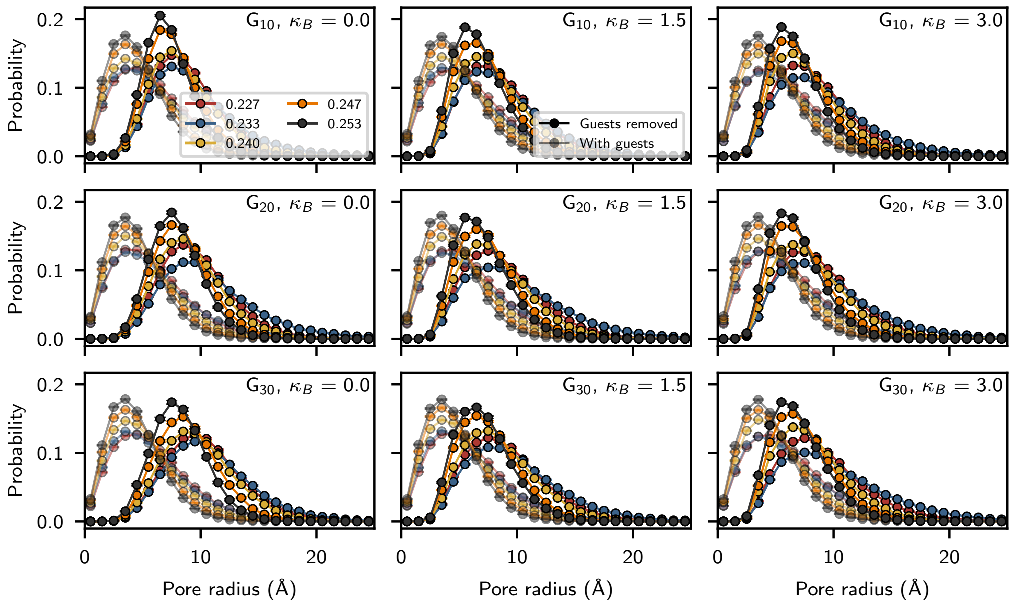


FIG. S25: Pore size distribution (PSD) measured from flux simulations of different guest molecules (length and stiffness) through host condensates with varying tyrosine fractions. PSD is measured with two variations: (i) with guest molecules included in the measurement (translucent curves), and (ii) with guest molecules removed (solid curves). In the latter, guest molecule positions are used as the probe points in the measurements.

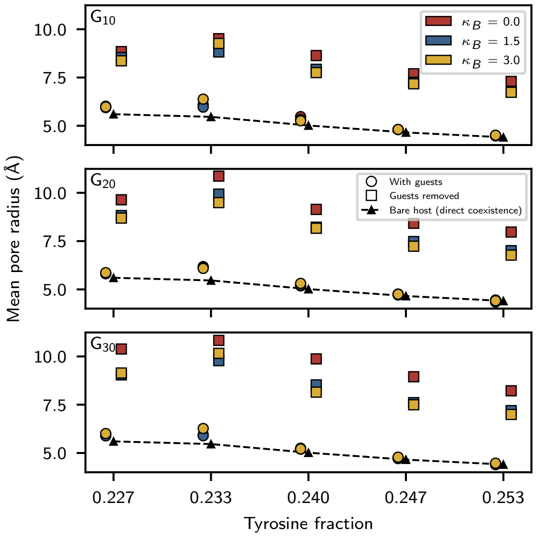


FIG. S26: Mean pore radius measured from flux simulations of different guests (varying length and stiffness) with host condensates of varying tyrosine fraction. Black triangles represent the mean pore sizes of the bare condensate, while the circles and squares come from measurements with and without the guest molecules in the PSD measurements [Fig. S25], respectively.
